# Mitochondrial outer membrane integrity regulates a ubiquitin-dependent NF-κB inflammatory response

**DOI:** 10.1101/2023.09.21.558776

**Authors:** Esmee Vringer, Joel S Riley, Annabel Black, Catherine Cloix, Sergio Lilla, Henning Walczak, Mads Gyrd-Hansen, Danny T Huang, Sara Zanivan, Stephen WG Tait

## Abstract

Mitochondria are often essential for apoptosis through mitochondrial outer membrane permeabilization (MOMP). This central event enables cytochrome *c* release leading to caspase activation and rapid cell death. Recently, MOMP has been shown to be inherently pro-inflammatory, for instance, by enabling mitochondrial DNA-dependent activation of cGAS-STING signalling. Alongside having emerging functions in health and disease, MOMP associated inflammation can also elicit anti-tumour immunity. Nonetheless, how MOMP triggers inflammation and how the cell counteracts this remains poorly defined. We find that upon MOMP, mitochondria are ubiquitylated in a promiscuous manner targeting proteins localised to both inner and outer mitochondrial membranes. Mitochondrial ubiquitylation serves to recruit the essential adaptor molecule, NEMO, leading to activation of pro-inflammatory NF-κB signalling. We find that disruption of mitochondrial outer membrane integrity through different means leads to engagement of a similar pro-inflammatory signalling platform. Thus, mitochondrial integrity directly controls inflammation, such that permeabilised mitochondria initiate NF-κB signalling. This event may be important for the various pathophysiological functions of MOMP-associated inflammation.

## Introduction

Apoptotic cell death is considered an immunosilent form of cell death, in line with it being the major type of homeostatic cell death. Mitochondrial outer membrane permeabilization (MOMP) is often essential to initiate apoptosis by enabling cytochrome *c* release, leading to rapid caspase activation and cell death (Bock & Tait, 2020). Nonetheless, upon a lethal stress, MOMP commits a cell to die regardless of caspase activation through so-called caspase-independent cell death (CICD). This is due to widespread MOMP causing a catastrophic loss in mitochondrial function (Lartigue *et al*, 2009).

Recent research has revealed that MOMP is inherently pro-inflammatory (Giampazolias *et al*, 2017; Marchi *et al*, 2022). For instance, mitochondrial DNA (mtDNA) is released from permeabilised mitochondria through BAX/BAK macropores, leading to activation of cGAS-STING signalling and a type I interferon response (McArthur *et al*, 2018; Riley *et al*, 2018). Importantly, while wholly dispensable for cell death, caspase activity serves to inhibit inflammation during mitochondrial apoptosis. Caspases inhibit inflammation in dying cells through multiple means, including direct cleavage of pro-inflammatory signalling proteins such as cGAS, inhibition of protein translation and promoting rapid removal of dying cells via the exposure of “eat-me” signals (McIlwain *et al*, 2013; Ning *et al*, 2019; Ravichandran, 2011).

By enhancing MOMP-induced inflammation through caspase-inhibition, we and others have shown that engaging CICD in tumour cells can lead to anti-tumour immunity dependent on cGAS-STING and NF-κB signalling in the dying cell (Giampazolias *et al*., 2017; Han *et al*, 2020). We also reported that MOMP can occur in a limited cohort of mitochondria – a phenomemon we termed minority MOMP– in the absence of cell death (Cao *et al*, 2022; Ichim *et al*, 2015). Minority MOMP can promote caspase-dependent DNA-damage. Intriguingly others have discovered that minority MOMP causes inflammation required for the restriction of bacteria. (Brokatzky *et al*, 2019). More recently, we have found that minority MOMP contributes to the inflammatory phenotype of senescent cells thereby directly bridging apoptotic signalling with senescence (Chapman et al, Res Square).

Thus, MOMP induced inflammation - alongside having physiological functions - represents a therapeutic target in cancer. Nonetheless, how MOMP elicits inflammation and how this is restrained remain poorly defined. Specifically, how permeabilised mitochondria are targeted for degradation – potentially limiting inflammation following MOMP, is not known. We initially set out to address this question, finding that upon MOMP, mitochondria are promiscuously ubiquitylated. Mitochondrial ubiquitylation has been shown to serve as signal for mitophagy, best evidenced in mitophagy promoted by the E3 ubiquitin ligase Parkin (Vargas *et al*, 2022). Surprisingly, upon MOMP, we find that autophagy is not essential for mitochondrial degradation. Upon further investigation, we found that MOMP-induced ubiquitylation of mitochondria serves as an inflammatory signal, recruiting the essential NF-κB signalling adaptor, NF-κB essential modulator (NEMO). In this way, mitochondrial outer membrane integrity dictates the initiation of an NF-κB inflammatory response, contributing to MOMP-induced inflammation.

## Results

### Permeabilised mitochondria are ubiquitylated and degraded independent of canonical autophagy

Following MOMP, autophagy targets permeabilised mitochondria for degradation and suppresses MOMP-induced inflammation (Colell *et al*, 2007; Lindqvist *et al*, 2018). Given this, our initial goal was to understand how MOMP triggers mitochondrial removal. To engage mitochondrial apoptosis, U2OS cells were treated a combination of BH3-mimetics, ABT-737 (inhibits BCL-2, BCL-xL and BCL-w) and S63845 (inhibits MCL-1) then analysed for cell viability by SYTOX Green exclusion and Incucyte live-cell imaging. Combined BH3-mimetic treatment in wild-type U2OS cells led to rapid cell death that was inhibited by co-treatment with pan-caspase inhibitor Q-VD-OPh or CRISPR-Cas9 mediated deletion of BAX and BAK, two proteins essential for MOMP, confirming engagement of mitochondrial apoptosis (**Supplemental Figures 1A** and **B**). Using this approach, we next assessed mitochondrial content in U2OS cells following MOMP under conditions of CICD by using the combination treatment of ABT-737, S63845 and Q-VD-OPh. Mitochondrial content was determined by western blot for mitochondrial proteins or via qPCR for mitochondrial DNA (**Figure 1A** and **1B)**. Reduction in cellular mitochondrial content was observed specifically following MOMP, as evidenced by a loss of mtDNA and mitochondrial protein content in a BAX/BAK dependent manner (**Figure 1A** and **1B**). Mitochondrial ubiquitylation is a well-established signal for autophagic removal of mitochondria, a process called mitophagy (Vargas *et al*., 2022). Therefore, we investigated whether mitochondria are ubiquitylated upon MOMP. SVEC4-10 murine endothelial cells were treated to undergo CICD and mitochondrial-enriched fractions were probed for ubiquitylation by western blot using a pan-ubiquitin antibody. Consistent with engagement of MOMP, SMAC (also called DIABLO) was depleted from the mitochondrial-enriched fraction upon CICD. Importantly, an extensive increase of protein ubiquitylation was detected in the mitochondria-enriched fraction specifically following MOMP (**Figure 1C**). Increased ubiquitylation was dependent upon MOMP since it was absent in BAX/BAK deficient cells (**Supplemental Figure 1C**). To corroborate these findings, U2OS EMPTY^CRISPR^ and BAX/BAK^CRISPR^ cells were immunostained following induction of CICD using a combination of anti-ubiquitin and mitochondrial COXIV antibodies. Upon CICD, ubiquitin localised with mitochondria in U2OS EMPTY^CRISPR^ cells but not in U2OS BAX/BAK^CRISPR^ cells (**Figure 1D** and **1E**), in line with the earlier mitochondrial fractionation experiment (**Supplemental Figure 1C**). To investigate whether inhibition of caspase activity was required for the ubiquitylation of mitochondria following MOMP, SVEC4-10 cells were treated with BH3-mimetics with or without the pan-caspase inhibitor Q-VD-OPh. Western blot analysis of mitochondria-enriched fractions demonstrated increased ubiquitylation irrespective of caspase inhibition (**Supplemental Figure 1D**).

**Figure 1.**
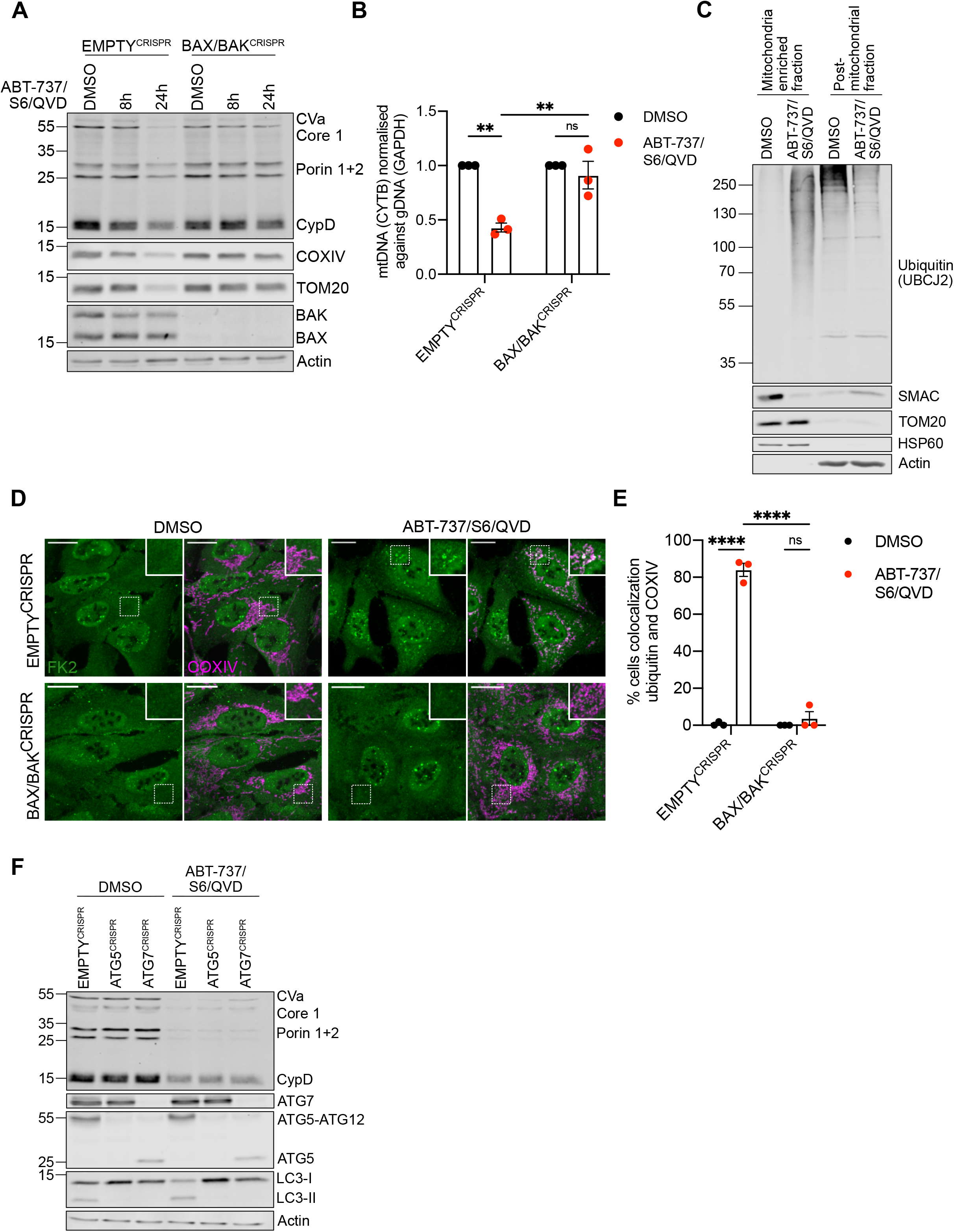
Mitochondrial depletion after MOMP does not require autophagy. A) U2OS EMPTY^CRISPR^ and BAX/BAK^CRISPR^ cells treated with 10 μΜ ABT-737, 2 μΜ S63845 and 20 μΜ Q-VD-OPh for 8 or 24 hours. Mitochondrial depletion was assessed by blotting for several mitochondrial proteins. Blot is representative of 3 independent experiments. B) U2OS EMPTY^CRISPR^ and BAX/BAK^CRISPR^ cells were treated with 10 μΜ ABT-737, 2 μΜ S62845 and 20 μΜ Q-VD-OPh for 24 hours. Graphs shows presence of mtDNA relative to gDNA in 3 independent experiments. C) SVEC4-10 cells treated for 1 hour with 10 μΜ ABT-737, 10 μΜ S63845 and 30 μΜ Q-VD-OPh. Mitochondria were isolated using dounce homogeniser. Lysates for blotted for ubiquitin (UBCJ2), SMAC, TOM20, HSP60 and actin. Blots are representative of 2 independent experiments. D) U2OS EMPTY^CRISPR^ and BAX/BAK^CRISPR^ cells treated for 3 hours with 10 μΜ ABT-737, 2 μΜ S63845 and 20 μΜ Q-VD-OPh. Cells were stained for ubiquitin (FK2) and mitochondrial COXIV. Images are representative of 3 independent experiments. Images are maximum projections of Z-stacks with a scale of 20 μm. E) Quantification of panel C showing the percentage of cells with mitochondrial localised ubiquitin puncti. F) U2OS EMPTY^CRISPR^, ATG5^CRISPR^ and ATG7^CRISPR^ expressing YFP-Parkin were treated with 10 μΜ ABT-737, 2 μΜ S63845 and 20 μΜ Q-VD-OPh for 24 hours. Mitochondrial depletion was assessed by blotting for several mitochondrial proteins. Blot is representative of 3 independent experiments. Statistics for all experiments were performed using two-way ANOVA with Tukey correction. * p < 0.05, ** p < 0.01, **** p < 0.0001.

Ubiquitylation can target organelles for autophagic degradation via recruitment of specific autophagy adaptor molecules (Vargas *et al*., 2022). Therefore, we investigated whether autophagy was required for degradation of mitochondria following MOMP by engaging CICD in U2OS cells deficient in ATG5 or ATG7, two proteins essential for canonical macroautophagy (Komatsu *et al*, 2005; Kuma *et al*, 2004). ATG5 and ATG7 loss, as well as functional autophagy deficiency, evident by an absence of lipidated LC3, was confirmed via western blot (**Figure 1F**). Surprisingly, treatment of cells with BH3-mimetics and caspase inhibitor led to a reduction of mitochondria (as determined by the loss of mitochondrial protein content) independent of autophagy (**Figure 1F**). These data demonstrate that upon MOMP, mitochondria are ubiquitylated and can be degraded in a manner that does not require canonical autophagy.

### Widespread mitochondrial protein ubiquitylation occurs upon MOMP

We next characterised mitochondrial protein ubiquitylation upon MOMP. Diglycine remnant proteomics can identify ubiquitylated proteins by the immunoprecipitation of modification diGly-motifs left on ubiquitylated proteins after trypsinisation (Xu *et al*, 2010). Using this method, we investigated the ubiquitylome of SVEC4-10 cells treated to undergo CICD. Mass spectrometry proteomic analysis revealed a significant change in the ubiquitylome of CICD treated SVEC4-10 cells compared to untreated (**Figure 2A**). Gene-ontology (GO) term analysis and manual curation of proteins using MitoCarta 3.0 (Rath *et al*, 2021) revealed that most peptides (approx. 80%) that gained a ubiquitin modification after MOMP were mitochondrially localised (**Figures 2B** – **D, Supplemental table 1**). Ubiquitylated mitochondrial proteins were not confined to one mitochondrial compartment, with broadly similar numbers of ubiquitylated proteins characterised as being localised to the mitochondrial outer membrane or mitochondrial inner membrane (**Figure 2B** and **Supplemental Table 1**). Notably, some proteins with increased ubiquitylation have been defined as being localised to the mitochondrial matrix, possibly reflecting mitochondrial inner membrane permeabilisation that we and others have reported previously (**Figure 2B** and **Supplemental Table 1**) (McArthur *et al*., 2018; Riley *et al*., 2018). These data demonstrate promiscuous ubiquitylation of mitochondrial proteins following MOMP.

**Figure 2.**
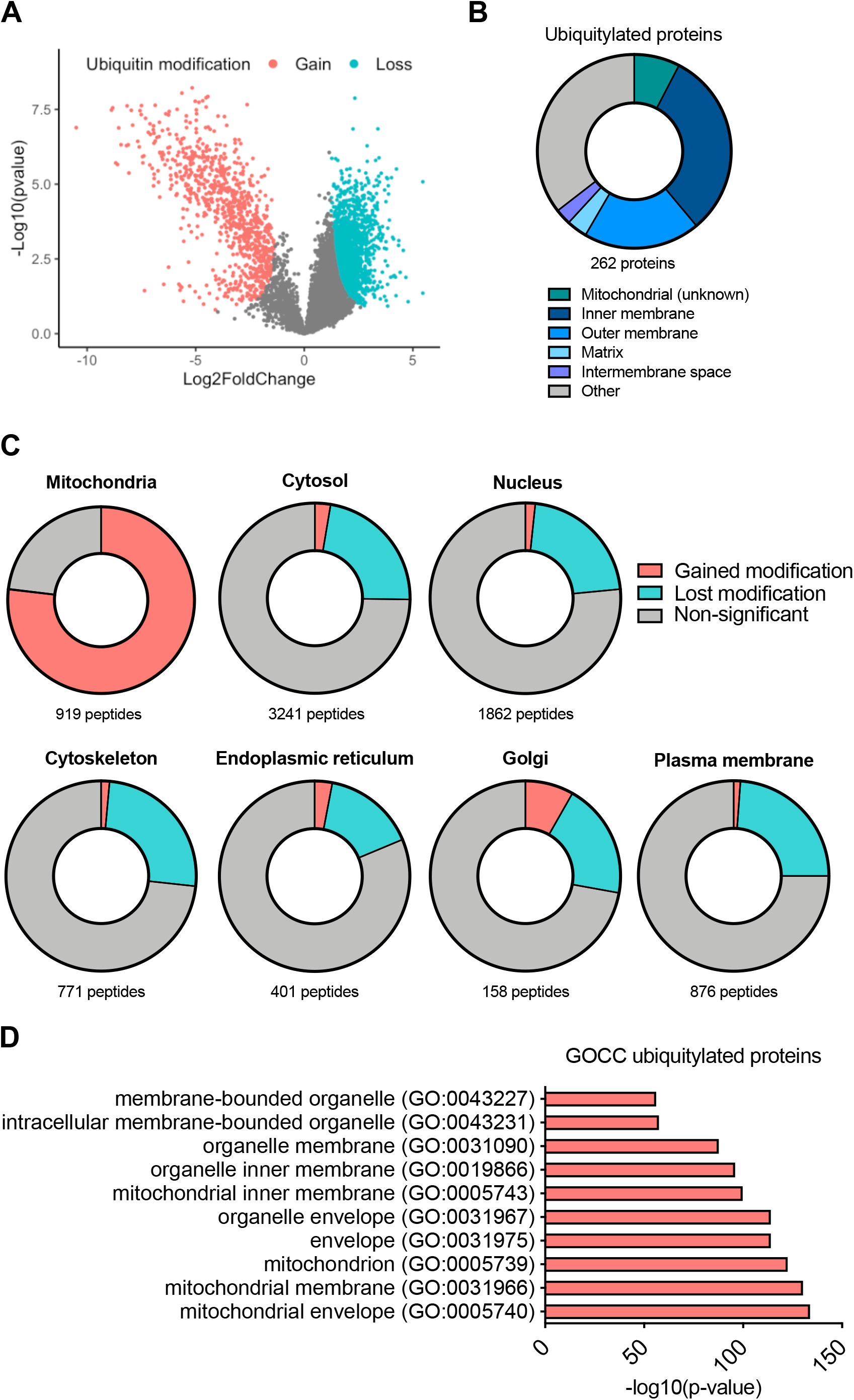
Widespread ubiquitylation of mitochondrial proteins after MOMP. A) Volcano plot of ubiquitylated proteins in SVEC4-10 cells treated for 3 hours with 10 μΜ ABT-737, 10 μΜ S63845 and 30 μΜ Q-VD-OPh. Experiment performed with 4 independent repeats. Significance (coloured dots) determined using Students T Test. Plot generated in R studio. B) Pie chart of ubiquitylated peptides categorised into mitochondrial compartments. Categorisation of peptides was performed using MitoCarta 3.0, UniProt and ProteinAtlas. C) Cellular distribution of all hits from the isolated ubiquitin remnant-containing peptides. Categorisation using MitoCarta3.0, UniProt and ProteinAtlas. D) GO-term cellular compartment analysis of proteins with increased ubiquitylation after MOMP. Graphs shows the top 10 most significant hits.

### Mitochondrial protein ubiquitylation encompasses K63-ubiquitin linkages

Protein ubiquitylation is highly complex with specific ubiquitin linkages conferring distinct biological functions. For instance, K48 ubiquitin linkages are typically associated with targeting proteins for proteasomal degradation, whereas K63 ubiquitylation has signalling functions (Komander & Rape, 2012). Given this, we investigated the type of ubiquitin linkages that MOMP triggers. SVEC4-10 cells were treated to undergo CICD, and the mitochondrial enriched fraction was blotted for K48-and K63-ubiquitin linkages using linkage specific antibodies (**Figure 3A**). This revealed an increase in K63-linked ubiquitin, but not K48-linked ubiquitin, in the mitochondrial fraction specifically during CICD. K63-linked ubiquitylation of mitochondria was also detected upon CICD by immunofluorescence (**Figure 3B** and **3C**). Finally, we made use of GFP-fused ubiquitin binding domains (UBDs) developed to specifically visualise K63 and linear M1 ubiquitin linkages (Hrdinka *et al*, 2016). Consistent with our previous data, extensive K63-linked ubiquitin was detected on mitochondria following CICD (**Figure 3D** and **3E**). In contrast, mitochondrial localisation of M1-specific UBDs was observed in a smaller percentage of cells analysed. These data reveal that upon MOMP, mitochondrial proteins are enriched in K63-ubiquitin linkages.

**Figure 3.**
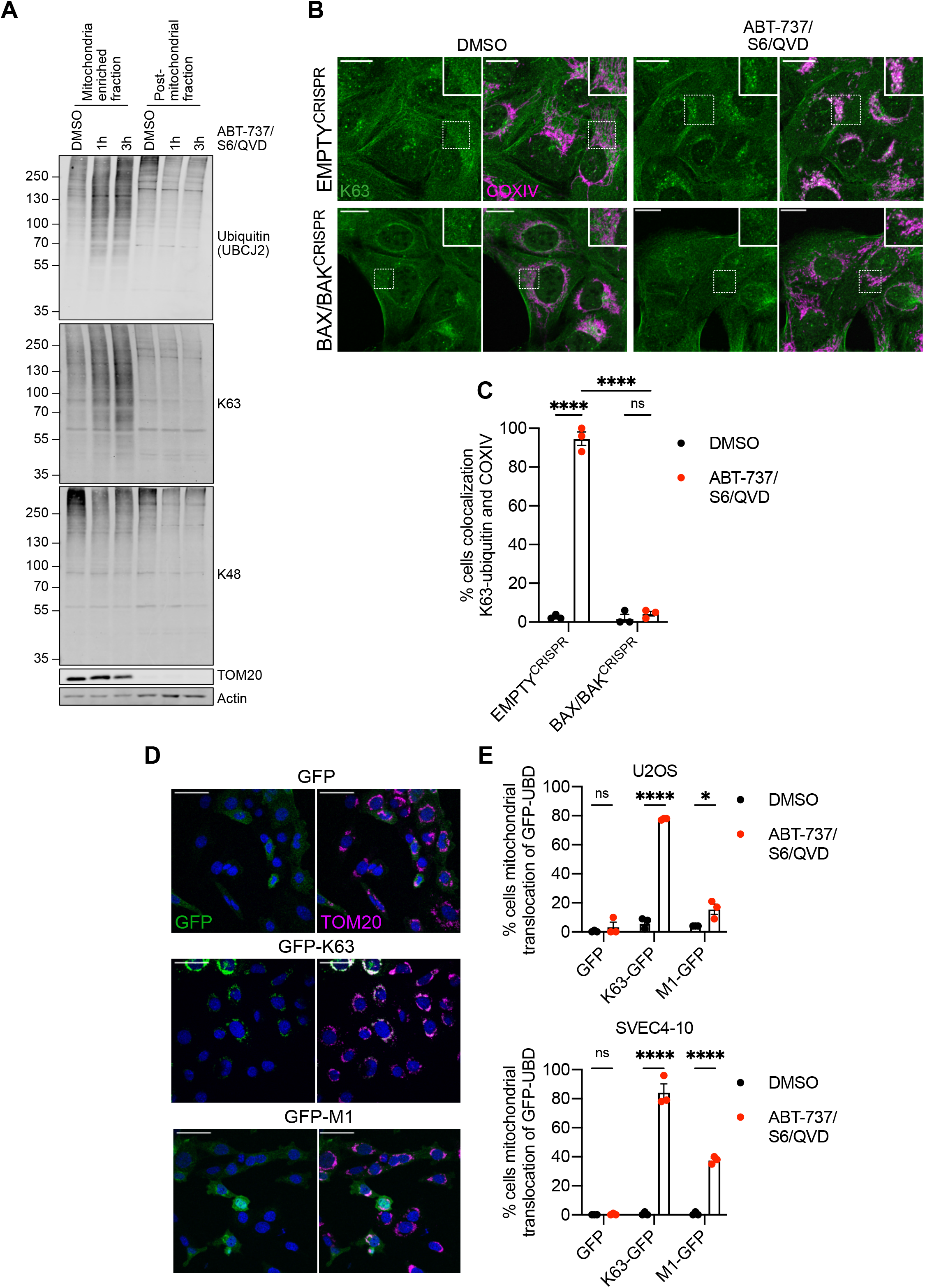
K63-linked ubiquitylation on mitochondria after MOMP. A) SVEC4-10 cells treated with for 1 or 3 hours with 10 μΜ ABT-737, 10 μΜ S63845 and 30 μΜ Q-VD-OPh. Mitochondria were isolated using digitonin fractionation buffer and antibodies against pan-ubiquitin (UBCJ2), K63- and K48-specific ubiquitin were used. Blots representative for 3 independent experiments. B) U2OS EMPTY^CRISPR^ and. BAX/BAK^CRISPR^ cells treated with 10 μΜ ABT-737, 2 μΜ S63845 and 20 μΜ Q-VD-OPh for 3 hours. Stained for K63-ubiquitin and COXIV. Images are maximum projections of Z-stacks with a scale of 20 μm and are representative of 3 independent experiments. C) Quantification of D showing the percentage of cells with mitochondrial localised K63-ubiquitin puncti. Statistics performed using two-way ANOVA with Tukey correction. D) SVEC4-10 cells expressing doxycycline-inducible K63 or M1-UBDs. Cells were treated for 1 hour with 10 μΜ ABT-737, 10 μΜ S63845 and 30 μΜ Q-VD-OPh. Images are representative of 3 independent experiments with a scale bar of 50 μm. E) Quantification of D showing the percentage of SVEC 4-10 cells with mitochondrial localised GFP-UBDs. Also includes the quantification of U2OS cells expressing doxycycline-inducible K63- or M1-UBDs treated for 3 hours with 10 μΜ ABT-737, 2 μΜ S63845 and 20 μΜ Q-VD-OPh. Statistics were performed using multiple unpaired t-tests. ** p < 0.01, *** p < 0.001. **** p < 0.0001.

### Mitochondrial ubiquitylation recruits the essential NF-κB adaptor NEMO promoting NF-κB activation

We next sought to understand potential biological functions of mitochondrial ubiquitylation following MOMP. Our previous data demonstrated that NF-κB is activated following MOMP, contributing to anti-tumorigenic effects of CICD (Giampazolias *et al*., 2017). This finding, coupled to the well-established connection between K63-ubiquiylation and inflammatory signalling (Madiraju *et al*, 2022), led us to investigate if mitochondrial ubiquitylation may be involved in NF-κB activation during CICD. NEMO, an essential adaptor protein in canonical NF-κB signalling, initiates NF-κB activity through ubiquitin binding. Given this, we examined the localisation of NF-κB essential modulator (NEMO) under conditions of MOMP, by expressing GFP-NEMO in U2OS cells (**Figures 4A** and **4B**). Importantly, robust mitochondrial translocation of GFP-NEMO occurred upon MOMP in a BAX/BAK-dependent manner. To investigate if ubiquitylation was required for mitochondrial recruitment of NEMO we used the E1 inhibitor TAK-243 to block ubiquitylation (Hyer *et al*, 2018). TAK-243 treatment effectively blocked mitochondrial ubiquitylation and mitochondrial recruitment of GFP-NEMO, (**Figure 4C-D** and **Supplemental Figure 2A**). In contrast, blocking the ubiquitin-like modification neddylation, using NAE1 inhibitor MLN4924 (Soucy *et al*, 2009), did not result in reduced ubiquitylation and GFP-NEMO translocation in SVEC4-10 cells (**Supplemental Figure 2B - D**).

**Figure 4.**
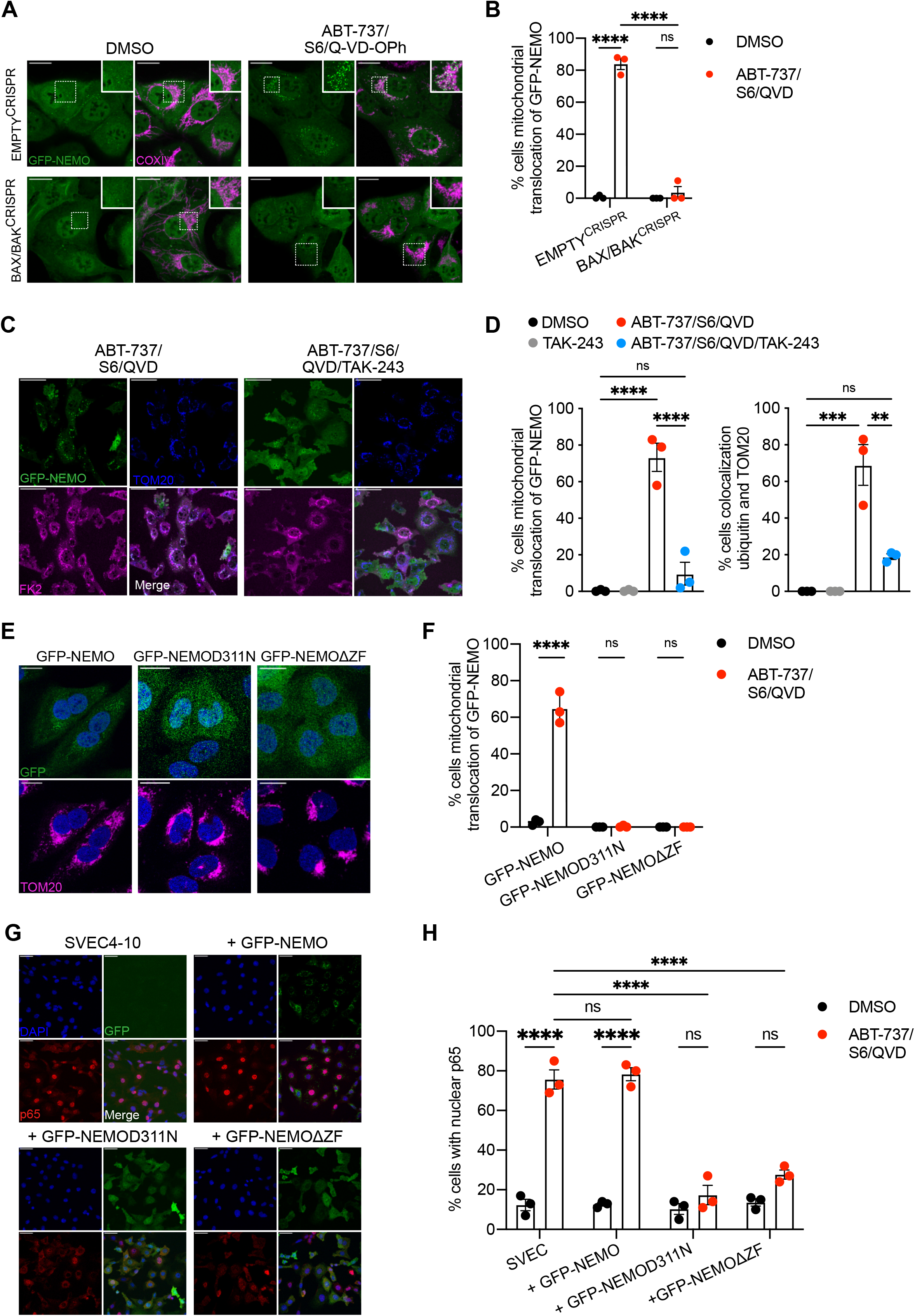
Ubiquitin-dependent recruitment of NEMO to mitochondria is essential for NF-κB activation after MOMP. A) U2OS EMPTY^CRISPR^ and BAX/BAK^CRISPR^ cells expressing GFP-NEMO were treated for 3 hours with 10 μΜ ABT-737, 2 μΜ S63845 and 20 μΜ Q-VD-OPh. Cells were immunostained for COXIV. Scale bar is 20 μm. Images are maximum projections of Z-stacks and are representative for 3 independent experiments. B) Quantification of A showing the percentage of cells with mitochondrial localised GFP-NEMO puncti. C) SVEC4-10 cells expressing GFP-NEMO were pre-treated for 1 hour with 2 μΜ TAK-243 followed by 1 hour treatment of 10 μΜ ABT-737, 10 μΜ S63845, 30 μΜ Q-VD-OPh with or without 2 μΜ TAK-243. Cells were immunostained for TOM20 and ubiquitin (FK2). Scale bar is 50 μm and images are representative for 3 independent experiments. D) Quantification of C showing the percentage of cells with mitochondrial localised GFP-NEMO and ubiquitin puncti. E) U2OS cells expressing GFP-NEMO, GFP-NEMOD311N or GFP-NEMOΔZF were treated for 3 hours with 10 μΜ ABT-737, 2 μΜ S63845 and 20 μΜ Q-VD-OPh. Cells were immunostained for TOM20 and DAPI. Scale bar is 20 μm and images are representative for 3 independent experiments. F) Quantification of E showing the percentage of cells with mitochondrial translocation of GFP-NEMO. G) Parental SVEC4-10 cells and SVEC4-10 cells expressing GFP-NEMO, GFP-NEMOD311N or GFP-NEMOΔZF were treated for 1 hour with 10 μΜ ABT-737, 10 μΜ S63845 and 30 μΜ Q-VD-OPh. Cells were immunostained for p65 and DAPI. Scale bar is 50 μm and images are representative for 3 independent experiments. H) Quantification of G showing the GFP+ cells with nuclear translocation of p65. Statistics are performed using two-way ANOVA with Tukey correction. ** p < 0.01, *** p < 0.001. **** p < 0.0001.

M1-ubiqutin linkages are often essential for NEMO activation, through binding the UBAN domain on NEMO (Rahighi *et al*, 2009). Additionally, the C-terminal zinc finger (ZF) domain of NEMO can enhance the binding of K63-ubiquitin chains to the UBAN (Cordier *et al*, 2009; Laplantine *et al*, 2009). To determine which domain(s) are required for ubiquitin-dependent recruitment after MOMP, mutant versions of NEMO disrupting the ability of the UBAN (D311N) or C-terminus (ΔZF) to bind ubiquitin were generated. Both non-ubiquitin binding mutants of NEMO failed to be recruited to the mitochondria after MOMP (**Figure 4E** and **4F**) suggesting that binding to K63-ubiquitin chains is required. To extend these findings, we made use of murine embryonic fibroblasts (MEFs) deficient in HOIP, the catalytic subunit of LUBAC E3 ligase complex required for M1-linked ubiquitylation (Peltzer *et al*, 2014). Importantly, mitochondrial recruitment of GFP-NEMO was not impaired in HOIP-deficient MEFs (**Supplemental Figures 3A** and **3B**). These data demonstrate that K63-linked ubiquitylation, but not M1-linked ubiquitylation, is required for NEMO recruitment to the mitochondria.

We next determined if mitochondrial recruitment of NEMO facilitates NF-κB activation. SVEC4-10 cells expressing wild-type and non-ubiquitin binding variants of GFP-NEMO (D311N and ΔZF) were treated to engage CICD and NF-κB activation was determined by nuclear NF-κB p65 translocation. In contrast to wild-type NEMO, both non-ubiquitin binding variants of NEMO significantly inhibited NF-κB activation, as determined by a reduction in nuclear p65 (**Figure 4G** and **4H**). Similar experiments were performed in SVEC4-10 cells treated with siRNA to deplete endogenous murine NEMO (**Supplemental Figures 3C - E**). Depletion of NEMO in parental SVEC4-10 cells completely abolished nuclear p65 translocation. As expected, this was rescued by ectopic expression of human GFP-NEMO. In contrast, expression of human GFP-NEMOD311N failed to restore NF-κB p65 nuclear translocation, agreeing with our previous data. These data support a model whereby K63-ubiquitylation of mitochondria following MOMP enables NEMO recruitment leading to NF-κB activation.

### Ubiquitin-dependent NF-κB activation after MOMP is independent of canonical mitochondrial E3 ligases

Through MOMP, our findings directly link mitochondrial integrity to mitochondrial ubiquitylation and pro-inflammatory signalling. We next sought to identify which ubiquitin E3 ligase(s) may be responsible for mitochondrial ubiquitylation. One candidate is the E3 ligase Parkin, since active Parkin causes widespread ubiquitylation of mitochondrial proteins (Sarraf *et al*, 2013). Nonetheless, both SVEC4-10 and U2OS cells used in our studies do not express detectable Parkin (**Figure 5A**), arguing that mitochondrial ubiquitylation following MOMP does not require Parkin. Parkin activity requires the kinase PINK1. Interestingly, PINK1 can also activate alternative E3 ligases such as ARIH1 (Villa *et al*, 2017). To investigate a potential role for PINK1, we generated PINK1^CRISPR^ SVEC4-10 cell lines (**Supplemental Figure 4A**). Confirming functional deletion, cells lacking PINK1 failed to recruit YFP-Parkin to mitochondria following CCCP treatment, in contrast to EMPTY^CRISPR^ cells (**Supplemental Figure 4B**). YFP-Parkin was not recruited to mitochondria after MOMP irrespective of PINK1 deletion (**Supplemental Figure 4B**). Importantly, recruitment of GFP-NEMO was not impaired by the deletion of PINK1 (**Figure 5B** and **5C**) indicating that PINK1 does not have a role in ubiquitin-dependent recruitment of NEMO after MOMP. The mitochondrial resident E3 ligases MUL1 (also called MAPL) and MARCH5 have roles in various cellular processes such as mitochondrial dynamics, protein import, cell death and inflammation (Braschi *et al*, 2009; Haschka *et al*, 2020; Phu *et al*, 2020; Shiiba *et al*, 2020). Single and double knockout MUL1^CRISPR^ and MARCH5^CRISPR^ SVEC4-10 cell lines were generated (**Supplemental Figure 4C** and **4D**), however no differences in IκΒα phosphorylation, a marker for NF-κB activation, were observed (**Figure 5D**). Moreover, no impact on mitochondrial ubiquitylation following MOMP was observed in SVEC4-10 MUL1MARCH5^CRISPR^ cells (**Figures 5E-F**). Interestingly, MARCH5 is degraded upon MOMP (**Figure 5D**) indicating that its ubiquitylation observed in the ubiquitin remnant proteomics study might be linked to proteasomal degradation (**Supplemental Table 1**). The E3 ligase XIAP was previously described for its involvement in the recruitment of endolysosomes through ubiquitylation of mitochondrial proteins after MOMP (Hamacher-Brady *et al*, 2014). XIAP^CRISPR^ SVEC4-10 cell lines were generated to validate the importance of XIAP in mitochondrial-driven inflammation (**Supplemental Figure 4E**). No differences were observed in expression of pro-inflammatory cytokines after MOMP (**Supplemental Figure 4F**), despite observing a small reduction in the percentage of cells with mitochondrial ubiquitylation and GFP-NEMO recruitment (**Figure 5G** and **5H**). Combined, these data demonstrate that established mitochondrial E3 ligases are not required for mitochondrial ubiquitylation following MOMP.

**Figure 5.**
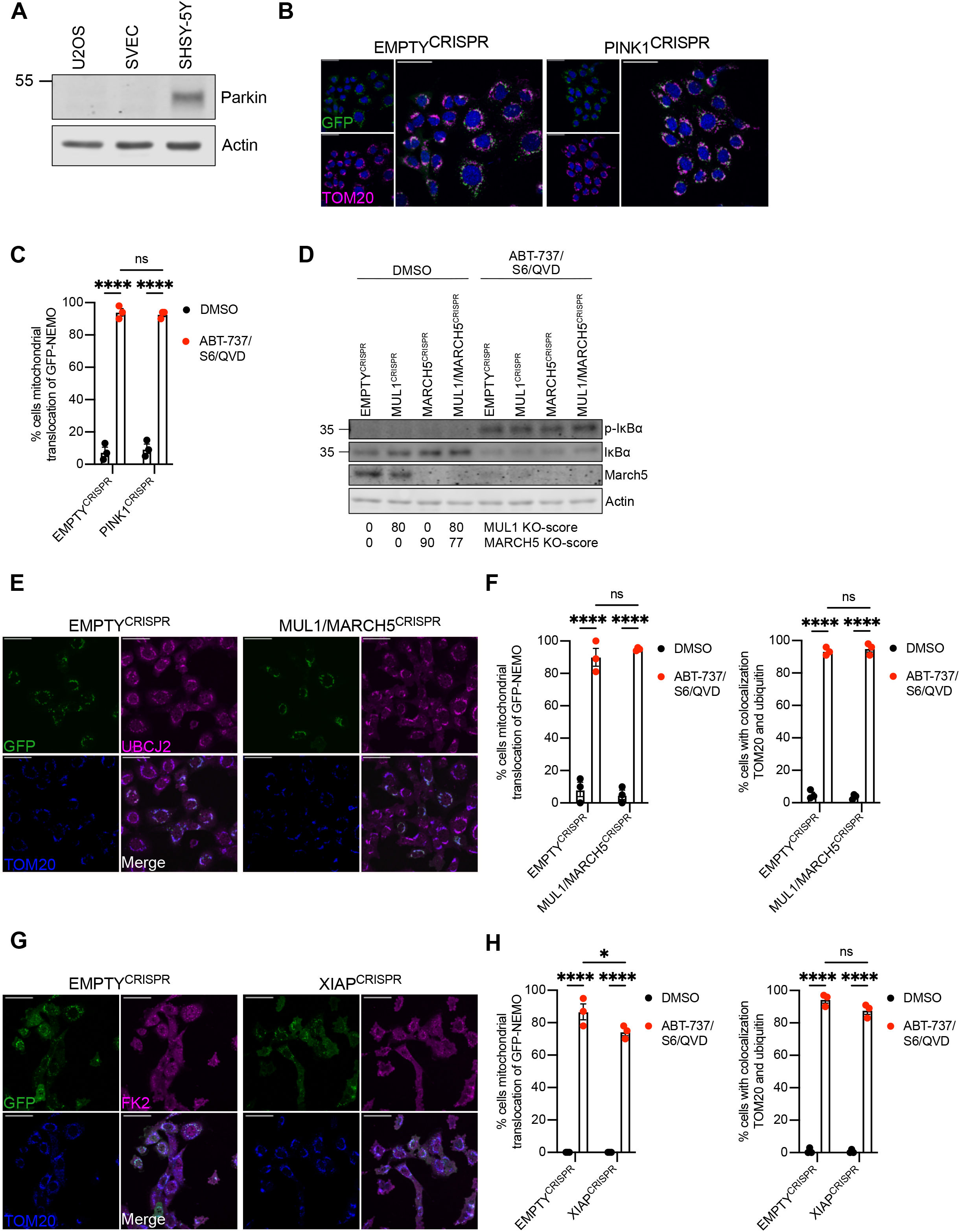
Ubiquitylation-induced inflammation after MOMP is independent of established mitochondrial E3 ligases. A) Lysates of U2OS, SVEC4-10 and SHSHY-5Y cells were blotted for Parkin and actin. B) SVEC4-10 EMPTY^CRISPR^ and PINK1^CRISPR^ cells expressing GFP-NEMO were treated for 1 hour with 10 μΜ ABT-737, 10 μΜ S63845 and 30 μΜ Q-VD-OPh. Cells were immunostained for TOM20. Images are representative for 3 independent experiments with a scale bar of 50 μm. C) Quantification of B showing the percentage of cells with mitochondrial translocation of GFP-NEMO. D) SVEC4-10 EMPTY^CRISPR^, MUL1^CRISPR^, MARCH5^CRISPR^ and MUL1/MARCH5^CRISPR^ treated for 3 hours with 10 μΜ ABT-737, 10 μΜ S63845 and 30 μΜ Q-VD-OPh. Lysates were blotted for p-ΙκΒα, ΙκΒα, MARCH5 and actin. Blots are representative of 3 independent experiments. KO-scores of MUL1 and MARCH5 are calculated via ICE analysis. E) SVEC4-10 EMPTY^CRISPR^ and MUL1/MARCH5^CRISPR^ cells expressing GFP-NEMO treated for 1 hour with 10 μΜ ABT-737, 10 μΜ S63845 and 30 μΜ Q-VD-OPh. Cells were immunostained for ubiquitin (UBCJ2) and TOM20. Images are representative for 3 independent experiments with a scale bar of 50 μm. F) Quantification of E showing the percentage of cells with mitochondrial localisation of GFP-NEMO and ubiquitin. G) SVEC4-10 EMPTY^CRISPR^ and XIAP^CRISPR^ cells expressing GFP-NEMO were treated with 10 μΜ ABT-737, 10 μΜ S63845 and 30 μΜ Q-VD-OPh for 1 hour. Cells were immunostained for ubiquitin (FK2) and TOM20 Images are representative for 3 independent experiments with a scale bar of 50 μm. H) Quantification of G showing the percentage of cells with mitochondrial localisation of GFP-NEMO and ubiquitin. Statistics were performed using two-way ANOVA with Tukey correction. **** p < 0.0001.

### Ubiquitin-dependent mitochondrial inflammation is regulated by mitochondrial outer membrane integrity

As discussed, mitochondrial apoptosis requires BAX and BAK activation leading to MOMP. We next sought to define if pro-inflammatory mitochondrial ubiquitylation was specific to mitochondrial apoptosis or initiated due to loss of mitochondrial integrity. For this purpose, we used the compound raptinal that can cause MOMP independent of BAX and BAK (Heimer *et al*, 2019; Palchaudhuri *et al*, 2015). In agreement, BAX/BAK deficient SVEC4-10 cells were protected against cell death induced by BH3-mimetics but remained sensitive to raptinal-induced cell death in a caspase-dependent manner (**Supplementary Figures 5A and B**). We next investigated GFP-NEMO translocation and mitochondrial ubiquitylation following raptinal treatment in BAX/BAK-deleted SVEC4-10 cells. Importantly, raptinal treatment led to robust mitochondrial ubiquitylation and GFP-NEMO translocation independently of BAX and BAK (**Figure 6A** and **6B**). Consistent with this, nuclear translocation of p65 was also observed in BAX/BAK deleted cells following raptinal treatment (**Figure 6C** and **6D**). Finally, increased transcription of NF-κB targets *Kc* and *Tnf* was detected following raptinal treatment in BAX/BAK-deleted SVEC4-10 cells (**Figure 6E**). Congruent with earlier findings, BH3-mimetic induced ubiquitylation, NEMO translocation and NF-κB activity required BAX and BAK (**Figure 6A-E**). These data demonstrate that loss of mitochondrial outer membrane integrity is sufficient to induce mitochondrial ubiquitylation leading to NEMO recruitment and an NF-κB dependent inflammatory response.

**Figure 6.**
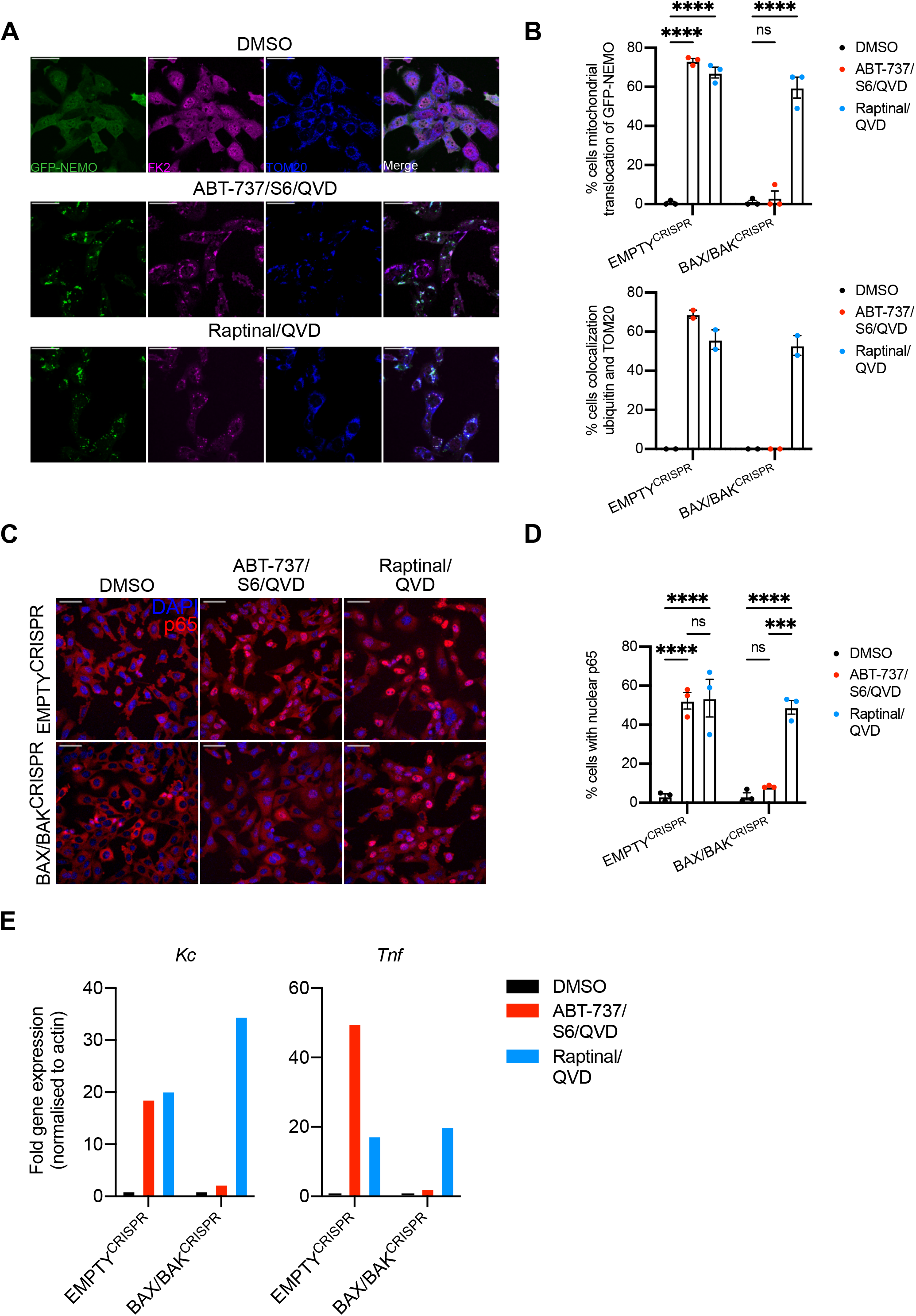
Mitochondrial ubiquitylation and inflammation occurs upon loss of mitochondrial outer membrane integrity. A) SVEC4-10 cells expressing GFP-NEMO were treated for 3 hours with 10 μΜ ABT-737, 10 μΜ S63845 and 30 μΜ Q-VD-OPh or 2.5 μΜ raptinal and 30 μΜ Q-VD-OPh. Cells were immunostained for ubiquitin (FK2) and TOM20. Images are representative of 2 independent experiments displayed with a 50 μm scale bar. B) Quantification of A showing the percentage of cells with mitochondrial localisation of GFP-NEMO and ubiquitin. Percentages of cells with mitochondrial localisation of GFP-NEMO was determined in 3 independent experiments, while mitochondrial localisation of ubiquitin was determined in 2 independent experiments. C) SVEC4-10 EMPTY^CRISPR^ and BAX/BAK^CRISPR^ cells were treated for 3 hours with 10 μΜ ABT-737, 10 μΜ S63845 and 30 μΜ Q-VD-OPh or 2.5 μΜ raptinal and 30 μΜ Q-VD-OPh. Cells were immunostained stained for p65 and DAPI. Images are representative of 3 independent experiments and are shown with a 50 μm scale bar. D) Quantification of B showing the percentage of cells with nuclear translocation of p65. E) SVEC4-10 cells treated for 3 hours with 10 μΜ ABT-737, 10 μΜ S63856 and 30 μΜ Q-VD-OPh or 2.5 μΜ raptinal and 30 μΜ Q-VD-OPh. Expression of *Kc*, *Tnf* and *Actin* were validated using RT-qPCR. Graphs are representative for 3 independent experiments. Statistics were performed using Dunnett correction. **** p < 0.0001.

## Discussion

We find that upon disruption of mitochondrial outer membrane integrity, mitochondria are promiscuously ubiquitylated; numerous proteins localising to both outer and inner mitochondrial membranes were found to be ubiquitylated. Investigating the functions of mitochondrial ubiquitylation, unexpectedly, we found that degradation of mitochondria could occur independently of canonical autophagy. We found that mitochondrial ubiquitylation directly promotes inflammatory NF-κB activation through mitochondrial recruitment of the adaptor molecule NEMO. These data connect mitochondrial outer membrane integrity to direct activation of NF-κB activity, contributing to the pro-inflammatory effects of MOMP.

Given the bacterial ancestry of mitochondria, our findings raise striking parallels with cell intrinsic responses to bacterial infection. For instance, ubiquitylation of intracellular *Salmonella* Typhimurium serves as a platform to initiate pro-inflammatory NF-κB signalling as an innate immune response (Noad *et al*, 2017; van Wijk *et al*, 2017)}. Notably, the mitochondrial inner membrane and bacterial membranes share similarities, for instance enrichment in cardiolipin (Vringer & Tait, 2022). We speculate that upon cytosolic exposure, the mitochondrial inner membrane may represent a damage-associated molecular pattern (DAMP) eliciting ubiquitylation, NEMO recruitment and inflammation. Nonetheless, distinct differences exist between NEMO recruitment leading to NF-κB activation on invading bacteria and permeabilised mitochondria. The most striking distinction is that, unlike bacteria, M1-linked ubiquitylation is not required for recruitment of NEMO to permeabilised mitochondria. This is best evidenced by mitochondrial recruitment of NEMO in cells deficient in HOIP, the catalytic subunit of LUBAC complex required for M1-linked ubiquitylation. Instead, NEMO recruitment to mitochondria appears dependent on its ability to bind K63-ubiquitylated proteins, indeed we observe extensive K63-linked (but not degradative K48-linked) mitochondrial ubiquitylation upon MOMP. Interestingly, a recent study has shown that mitochondria amplify TNF induced NF-κB signalling (Wu *et al*, 2022). In this paradigm, the mitochondrial outer membrane serves as a platform for LUBAC activity enhancing linear M1-linked ubiquitylation of NEMO. Together with our data, this positions mitochondria in different contexts as both initiators and amplifiers of NF-κB dependent signalling.

Mechanistic questions remain – not least the identity of the ubiquitin E3 ligase(s) required for MOMP induced ubiquitylation. Our data argues against key roles for PINK1/Parkin, XIAP or resident mitochondrial ubiquitin ligases such as MARCH5 and MUL1. Secondly, what properties of permeabilised mitochondria that initiates ubiquitylation remains unknown. Importantly, our data shows that mitochondrial ubiquitylation occurs upon loss of mitochondrial outer membrane integrity, independent of how this is achieved. This is best evidenced by MOMP engaged by either BAX/BAK or using the drug raptinal (in BAX/BAK null cells) both cause mitochondrial ubiquitylation, NEMO recruitment and NF-κB activation. While speculative, possibly proteins located on the inner mitochondrial membrane when exposed to the cytosol recruit and activate cytosolic ubiquitin ligases.

Our initial premise for this study stemmed from the hypothesis that mitochondrial ubiquitylation may serve as targeting signal for mitophagy, akin to PINK1/Parkin mediated mitophagy. Surprisingly, we found that mitochondrial degradation occurred in cells deficient in canonical autophagy. While this doesn’t negate a role for autophagy in promoting removal of permeabilised mitochondria, it demonstrates that autophagy not essential. Along these lines, we and others have previously found that upon MOMP, the mitochondrial outer membrane can be completely lost leaving what we called mito-corpses (Ader *et al*, 2019; Riley *et al*., 2018). Whether autophagy independent degradation of permeabilised mitochondria occurs in a regulated manner remains an open question.

In summary, our data reveals a novel direct connection between mitochondrial function and engagement of inflammation, where disruption of mitochondrial integrity initiates pro-inflammatory NF-κB signalling through extensive ubiquitylation and NEMO recruitment. Given the numerous emerging functions of MOMP induced inflammation, ranging from senescence to innate and anti-tumour immunity, basic understanding of this process may reveal new therapeutic opportunities.

## Materials and Methods

### Cell culture and chemicals

HEK293FT, SVEC4-10, MEFs and U2OS cells were cultured in high glucose DMEM supplemented with 10% FBS (Gibco #10438026), 2 mM glutamine (Gibco #25030081) and 1 mM sodium pyruvate (Gibco #11360070). Cells were cultured in 21% O2 and 5% CO2 at 37 °C. MEF *Tnf^-/-^ Hoip^+/+^* and MEF *Tnf^-/-^ Hoip^-/-^* cell lines have been described before (Peltzer *et al*., 2014). SVEC4-10 cells were purchased from ATCC. All cell lines were routinely tested for mycoplasma.

The following chemicals were used in this study: ABT-737 (APEXBIO #A8193), S63845 (Chemgood #C-1370), Q-VD-OPh (AdooQ Bioscience #A14915-25), Doxycycline hyclate (Sigma-Aldrich #D9891), TAK-243 (MedChemExpress #HY-100487), MLN4924 (Selleck Chemical #S7109), MG-132 (Selleck Chemical #S2619), and raptinal (Millipore Sigma #SML1745).

### Viral transfection

Overexpression and CRISPR cells lines were generated using lenti- or retroviral infection. For lentiviral transfections 1 μg VSVG (Addgene #8454) and 1.86 μg psPAX2 (Addgene #12260) were used. For retroviral transfections 1 μg VSVG and 1.86 μg HIV gag-pol (Addgene #14887) were used. For both transfections 5 μg of plasmid was used. HEK293FTs were transfected using lipofectamine 2000 or lipofectamine 3000 according to manufacturer’s instructions. After two days, virus-containing media was removed from the HEK293FTs and supplemented with 10 μg/ml polybrene before transferring to target cells. Two days after infection cells were selected using 2 μg/mL puromycin, 10 μg/mL blasticidin or 800 μg/mL neomycin. Some U2OS and SVEC4-10 lines expressing GFP were sorted for GFP expression instead of antibiotic selection.

The M6PblastGFP-NEMO, PMD-OGP and PMD-VSVG plasmids were gifted by Felix Randow. The pLenti-CMV-TetRepressor, pDestination-eGFP-NES, pDestination-eGFP-SK63-NES and pDestination-eGFP-NCM1-NES plasmids were gifted by Mads Gyrd-Hansen. CRISPR cell lines were generated using lentiCRISPRv1 or lentiCRISPRv2 vector (Addgene #52961) containing puromycin, blasticidin and neomycin resistance.

Human *BAK* 5’- GCCATGCTGGTAGACGTGTA −3’

Human *BAX* 5’- AGTAGAAAAGGGCGACAACC −3’

Human *ATG5* 5’- AAGAGTAAGTTATTTGACGT −3’

Human *ATG7* 5’- GAAGCTGAACGAGTATCGGC −3’

Mouse *Bak* 5’- GCGCTACGACACAGAGTTCC −3’

Mouse *Bax* 5’- CAACTTCAACTGGGGCCGCG −3’

Mouse *March5* 5’- AAGTACTCGGCGTTGCACTG −3’

Mouse *Mul1* 5’- TATATGGAGTACAGTACGG −3’

Mouse *Pink1* 5’- CTGATCGAGGAGAAGCAGG −3’

Mouse *Xiap* 5’- CATCAACATTGGCGCGAGCT −3’

### Generation of GFP-NEMOD311N and GFP-NEMOΔZF

GFP-NEMOD311N and GFP-NEMOΔZF were cloned into a pBABE-puro vector using EcoRI and BamHI restriction sites. GFP-NEMOD311N was cloned into the pBABE vector using Gibson assembly. NEMOD311N was obtained by PCR of pGEX- NEMOD311N (Addgene #11968). GFP was obtained by PCR of a GFP-containing plasmid. GFP-NEMOΔZF was obtained by PCR of the M6P-GFP-NEMO plasmid (gifted by Felix Randow), thereby removing the last 25 amino acids of wildtype NEMO.

GFP 5’- tctaggcgccggccggatccATGGTGAGCAAGGGCGAG −3’

GFP 3’- cagaaccaccaccaccCTTGTACAGCTCGTCCATGC −5’

NEMOD311N 5’-ctgtacaagggtggtggtggttctggtggtggtggttctAATAGGCACCTCTGGAAG −3’

NEMOD311N 3’- accactgtgctggcgaattcCTACTCAATGCACTCCATG −5’

GFP-NEMOΔZF 5’- TAAGCA GGATTCATGGTGAGCAAGGGCGAGGAG −3’

GFP-NEMOΔZF 3’- TGCTTA GAATTC CTAGTCAGGTGGCTCCTCGGGGG −5’

### ICE analysis for CRISPR

Genomic DNA was isolated from the empty vector and CRISPR cells. A PCR reaction for the CRISPR’ed region was set-up using Phusion DNA polymerase according to manufacturer’s instructions. The reactions were run on 2% agarose gel and bands of correct size were isolated and purified using the GeneJET Gel Extraction kit. Samples were sequenced by Eurofins genomics and analysed using ICE software by Synthego.

Mouse *March5* 5’- TCCTGGCCTGAAGGGTAGGGGA −3’

Mouse *March5* 3’- CCTCTTCCTTCCCCCACCCCAA −5’

Mouse *Mul1* 5’- GGGTCGCAGGTGATTTCGAGGC −3’

Mouse *Mul1* 3’- CACGTTGGAATCACCCCTGCCT −5’

Mouse *Pink1* 5’- TGTTGTTGTCCCAGACGTTTGT −3’

Mouse *Pink1* 3’- TAAATTGCCCAATCACGGCTCA −5’

### Knockdown using siRNA

SVEC4-10 cells were transfected with 20 nM siGENOME *Nemo* SMARTpool (Horizon Discovery #M-040796-01-0005) or siGENOME non-targeting control (Dharmacon #D0012061305) using lipofectamine RNAiMAX (Invitrogen #1377075). Experiments were performed 48 hours after transfection.

### RT-qPCR

RNA was isolated using the GeneJET RNA isolation kit (Thermo Fisher Scientific #K0732) according to manufacturer’s instructions. Genomic DNA was digested using an on-column DNase step (Sigma-Aldrich #04716728001) for 15 minutes. RNA was converted into cDNA using the High Capacity cDNA Reverse Transcriptase kit (Thermo Fisher Scientific #43-688-13) according to manufacturer’s instructions. cDNA was synthesised according to the following steps: 25 °C for 10 minutes, 37 °C for 120 minutes and 85 °C for 5 minutes.

RT-qPCR was performed by using the Brilliant III SYBR® Green QPCR Master Mix (Agilent #600882) or DyNAmo HS SYBR Green (Thermo Scientific #F410L) and the QuantStudio 3. The following RT-qPCR cycling parameters were used: initial denaturation on 95 °C for 10 minutes, 40 cycles of 95 °C for 20 seconds, 57 °C for 30 seconds and 72 °C for 30 seconds, finished by a dissociation step 65-95 °C (0.5

°C/second). Samples were run in technical triplicates. Fold change expression was determined using the 2^-ΔΔCT^ method.

### cDNA

Mouse *Actin* 5’- CTAAGGCCAACCGTGAAAAG −3’

Mouse *Actin* 3’- ACCAGAGGCATACAGGGACA −5’

Mouse *Tnf-α* 5’- GTCCCCAAAGGGATGAGAAG −3’

Mouse *Tnf-α* 3’- CACTTGGTGGTTTGCTACGAC −5’

Mouse *Kc* 5’- GGCTGGGATTCACCTCAAGAA −3’

Mouse *Kc* 3’- GAGTGTGGCTATGACTTCGGTT −5’

Mouse *Ccl5* 5’- CTGCTGCTTTGCCTACCTCT −3’

Mouse *Ccl5* 3’- CGAGTGACAAACACGACTGC −5’

### DNA

Human *CYTB* 5’- GCCTGCCTGATCCTCCAAAT −3’

Human *CYTB* 3’- AAGGTAGCGGATGATTCAGCC −5’

Human *GAPDH* 5’- TGGGGACTGGCTTTCCCATAA −3’

Human *GAPDH* 5’- CACATCACCCCTCTACCTCC −3’

### Western blotting

Cells were lysed in RIPA buffer (10 mM Tris-HCl (pH 7.4), 150 mM NaCl, 1.2 mM EDTA, 1% Triton X-100 and 0.1% SDS supplemented with cOmplete protease inhibitors) and proteins were isolated by maximal centrifugation (15,000 rpm) for 10 minutes. Lysates were loaded on 8, 10 or 12% gels and transferred onto nitrocellulose membranes. The membranes were blocked with 5% milk or BSA in TBS for 1 hour followed by overnight incubation of 1:1000 dilution of primary antibodies in 5% milk or BSA in TBST. The next day membranes were incubated with a 1:10,000 dilution of secondary antibodies for 1 hour and imaged on the Li-cor CLx. Primary antibodies used are actin (Sigma #A4700), ATG5 (CST #8540), ATG7 (CST #8558), LC3 (CST #2775), BAK (CST #12105), BAX (CST #2772), NEMO (Abcam #178872), FK2 (ENZO #BML-PW8810-0100), HSP60 (Santa Cruz #sc-13115), p-ΙκΒα (CST #2859), ΙκΒα (CST #4814), K48-ubiquitin (CST #8081), K63-ubiquitin (Merck #05-1308), NEDD8 (Abcam #AB81264), Parkin (Santa Cruz #sc−32282), SMAC (Abcam #AB32023), TOMM20 (Proteintech #11082-1-AP), UBCJ2 (ENZO #ENZ-ABS840-0100), Membrane Integrity Antibody cocktail (Abcam #ab110414), COXIV (CST #11967), MARCH5 (EMD Millipore #06-1036) and XIAP (BD #610716). Secondary antibodies used are goat anti-rabbit IgG (H+L) Alexa Fluor Plus 800 (Invitrogen #A32735), goat anti-mouse IgG (H+L) Alexa Fluor 680 (Invitrogen #A21057) and goat anti-mouse IgG (H+L) Dylight 800 (Invitrogen #SA535521).

### Mitochondrial isolation using digitonin

Cells were lysed in digitonin lysis buffer (0.25 M sucrose, 700 mM Tris-HCl pH 8 and 100 μg/mL digitonin) for 10 minutes on ice. The mitochondrial fraction was pelleted at 3000*g* for 5 minutes. Supernatant was stored as the non-mitochondrial fraction, the pellet was resuspended in RIPA lysis buffer and stored on ice for 20 minutes followed by centrifugation for 10 minutes at maximum speed (15,000 rpm). Supernatant was taken as mitochondrial fraction.

### Mitochondrial isolation using douce homogeniser

Cells were resuspended in mitochondrial isolation buffer (200 mM mannitol, 70 mM sucrose, 10 mM HEPES, 1 mM EGTA, pH 7.0, cOmplete protease inhibitor). After resuspension cells were homogenised using the dounce tissue grinder by performing 50 strokes up/down manually and centrifuged at 2000 rpm for 5 minutes. Supernatant was collected and pellet was resuspended in mitochondrial isolation buffer and spun down as previously described. Supernatant from both spins were combined and spun down at 9000 rpm for 5 minutes. The supernatant was kept as non-mitochondrial fraction. The pellet was resuspended in RIPA buffer and placed on ice for 20 minutes followed by centrifugation at maximum speed (15,000 rpm) for 10 minutes. Supernatant was kept as mitochondrial fraction.

### Immunofluorescent staining

Cells were fixed using 4% PFA for 15 minutes, followed by a 15 minutes permeabilization step using 0.2% Triton X-100. Samples were blocked using 2% BSA in PBS for 1 hour and incubated with primary antibody in 2% BSA overnight. The following day samples were incubated with secondary antibody in 2% BSA. Primary antibodies used are COXIV (CST #11967 and #4850), cytochrome *c* (BD #556432), FK2 (ENZO #BML-PW8810-0100), HSP60 (Santa Cruz #sc-13115), K63-ubiquitin (Merck #05-1308), p65 (CST #8242), TOMM20 (CST #42406 and Proteintech #11082-1-AP) and UBCJ2 (ENZO #ENZ-ABS840-0100). Secondary antibodies used are Alexa Fluor 488 goat anti-rabbit IgG (H+L) (Invitrogen #A11034), Alexa Fluor 488 goat anti-mouse IgG (H+L) (Invitrogen #A11029), Alexa Fluor 568 goat anti-rabbit IgG (H+L) (Invitrogen #A11011), Alexa Fluor 568 goat anti-mouse IgG (H+L) (Invitrogen #A11004), Alexa Fluor 647 goat anti-rabbit IgG (H+L) (Invitrogen #A21245) and Alexa Fluor 647 goat anti-mouse IgG (H+L) (Invitrogen #A21236). Coverslips were mounted using Vectashield with or without DAPI.

### Confocal microscopy

Cells were imaged using the Nikon A1R confocal microscope using all four lasers (405 nm, 488 nm, 561 nm and 638 nm) and images are acquired using sequential scanning. For p65 staining the 40x NA 1.30 oil-immersion objective was used, while the 60x 1.40 NA oil-immersion objective was used to determine ubiquitin, GFP-NEMO and YFP-Parkin puncta. Images were analysed using ImageJ version 2.1.0/1.53c and cells were counted using the cell counter plugin. Images may be displayed using pseudocolours.

### Cell death assays using Incucyte

Cell death assays were performed using Incucyte ZOOM from Sartorius. Cell death was measured by Sytox Green inclusion (Thermo Fisher Scientific #S7020). Images were taken every hour with a 10x objective. Starting confluency was used to normalisation.

### Isolation of peptides containing ubiquitin remnants

Peptides containing ubiquitin remnant motifs were isolated using the PTMScan® Ubiquitin Remnant Motif (K--GG) Kit (CST #5562) according to manufacturers’ instructions. Isolation of ubiquitin remnants was performed on 4 independent repeats for both conditions (4.4 mg protein per sample). Cellular localisation of proteins was determined using Uniprot and Proteinatlas. Mitochondrial localisation was determined using MitoCarta 3.0. GO enrichment analysis was performed using PANTHER classification system.

### Mass spectrometry

Peptides were separated by nanoscale C18 reverse-phase liquid chromatography using an EASY-nLC II 1200 (Thermo Scientific) coupled to an Orbitrap Fusion Lumos mass spectrometer (Thermo Scientific). Elution was performed at a flow rate of 300 nL/min using a binary gradient, into a 50 cm fused silica emitter (New Objective) packed in-house with ReproSil-Pur C18-AQ, 1.9 μm resin (Dr Maisch GmbH), for a total duration of 135 minutes. Packed emitter was kept at 50 °C by column oven (Sonation) integration into the nanoelectrospray ion source (Thermo Scientific). Eluting peptides were electrosprayed into the mass spectrometer using a nanoelectrospray ion source. To decrease air contaminants signal level an Active Background Ion Reduction Device (EDI Source Solutions) was used. Data acquisition was performed using Xcalibur software (Thermo Scientific). A full scan over mass range of 350-1400 m/z was acquired at 120,000 resolution at 200 m/z. Higher energy collision dissociation fragmentation was performed on the 15 most intense ions, and peptide fragments generated were analysed in the Orbitrap at 15,000 resolution.

The MS Raw data were processed using MaxQuant software version 1.6.3.3 and searched with Andromeda search engine (Cox *et al*, 2011) querying SwissProt (Consortium 2019) Mus musculus (20/06/2016; 57,258 entries). First and main searched were performed with precursor mass tolerances of 20 ppm and 4.5 ppm, respectively, and MS/MS tolerance of 20 ppm. The minimum peptide length was set to six amino acids and specificity for trypsin cleavage was required. Methionine oxidation, N-terminal acetylation and di-Gly-lysine were specified as variable modifications, whereas cysteine carbamidomethylation was set as fixed modification. The peptide, protein, and site false discovery rate (FDR) was set to 1%. All MaxQuant outputs were analysed with Perseus software version 1.6.2.3 (Tyanova *et al*, 2016). The MaxQuant output GlyGly (K)sites.txt file was use for quantification of Ubiquitylated peptides. From the GlyGly (K)Sites.txt file, Reverse and Potential Contaminant flagged peptides (defined as MaxQuant output) were removed. To determine significanly changing ubiquitylated peptides a Student t-test with a 1% FDR (permutation-based) was applied using the peptide intensities included in the GlyGly (K)Sites table. Missing values were imputated separately for each column (width 0.3, down shift 1.4). Only ubiquitylated peptides having: “score diff” greater than 5, a localisation probability higher than 0.75, and are robustly quantified in three out of four replicate experiments were included in the analysis.

## Data availability

The raw files and the MaxQuant search results files have been deposited to the ProteomeXchange Consortium (Deutsch *et al*, 2020) via the PRIDE partner repository (Perez-Riverol *et al*, 2022) with the dataset identifier PXD040192. Data are available via ProteomeXchange with identifier PXD040192.

For reviewer access: **Username:**

**Password:** YDEFnxY5

## Statistics

Statistics was performed using Prism 9. All data represent mean ± standard error of the mean (SEM) unless indicated differently.

* p < 0.05, ** p < 0.01, *** p < 0.001, **** p < 0.0001

## Acknowledgements

This research was supported by funding from the Cancer Research UK (A20145; S.W.G.T.)(A29256; D.T.H.). Stand Up to Cancer campaign for CRUK A29800 to S.Z.; A31287 to CRUK Beatson Institute, A18076 to CRUK Glasgow Centre, A17196 to CRUK Beatson Institute Advanced Technology Facilities. H.W. is supported by an Alexander von Humboldt Foundation Professorship Award, a Cancer Research UK Programme Grant (A27323), a Wellcome Trust Investigator Award (214342/Z/18/Z), a Medical Research Council Grant (MR/S00811X/1) and three collaborative research centre grants funded by Deutsche Forschungsgemeinschaft (DFG, German Research Foundation): SFB1399, Project C06; SFB1530-455784452, Project A03; and SFB1403–414786233. M.G-H. is supported by the LEO foundation and The Novo Nordisk Foundation (NNF20OC0059392). We thank Rosalie Heilig, Asma Ahmed and Catherine Winchester for critical reading of the manuscript. The authors declare that they have no conflict of interest

---

**Supplemental Figure 1.**
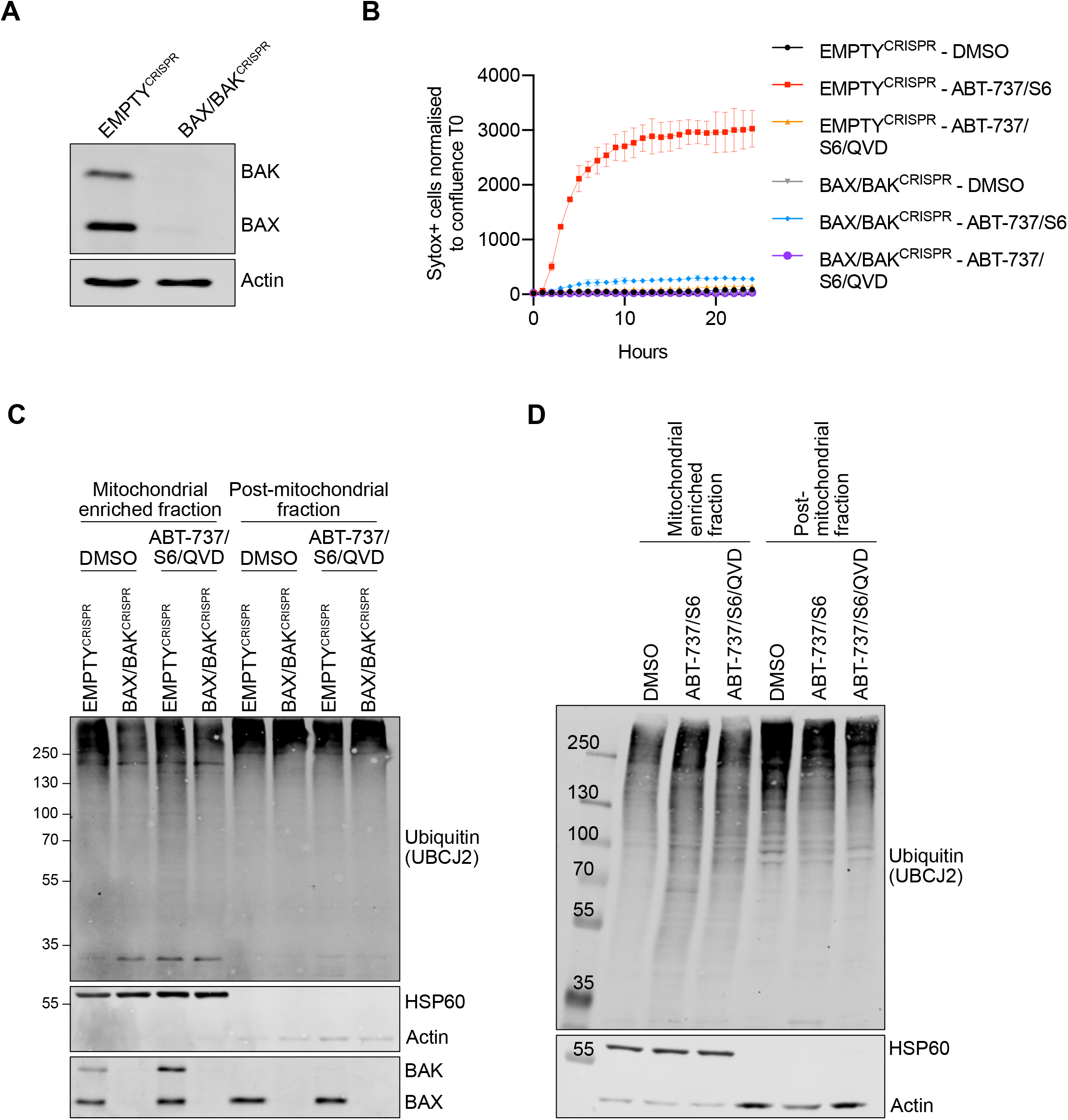
Ubiquitylation of mitochondria is dependent on MOMP by BAX/BAK pores, but independent of caspase activity. A) Lysates from U2OS EMPTY^CRISPR^ and BAX/BAK^CRISPR^ cells were blotted for BAX, BAK and Actin. B) U2OS EMPTY^CRISPR^ and BAX/BAK^CRISPR^ cells were treated with 10 μΜ ABT-737, 2 μΜ S63845 with or without 20 μΜ Q-VD-OPh. Cell death was monitored using Sytox Green inclusion normalised to starting confluence. Graph is representative for 3 independent experiments. C) SVEC4-10 EMPTY^CRISPR^ and BAX/BAK^CRISPR^ cells were treated for 1 hour with 10 μΜ ABT-737, 10 μΜ S63845 and 30 μΜ Q-VD-OPh. Mitochondria were isolated using digitonin fractionation buffer and blotted for ubiquitin (UBCJ2), BAX, BAK, HSP60 and actin. Blots are representative for 3 independent experiments. D) SVEC4-10 cells were treated for 1 hour with 10 μΜ ABT-737, 10 μΜ S63845 with or without 30 μΜ Q-VD-OPh. Mitochondria were isolated using digitonin fractionation buffer and lysates were blotted for ubiquitin (UBCJ2), HSP60 and actin. Blots are representative for 3 independent experiments.

**Supplemental Figure 2.**
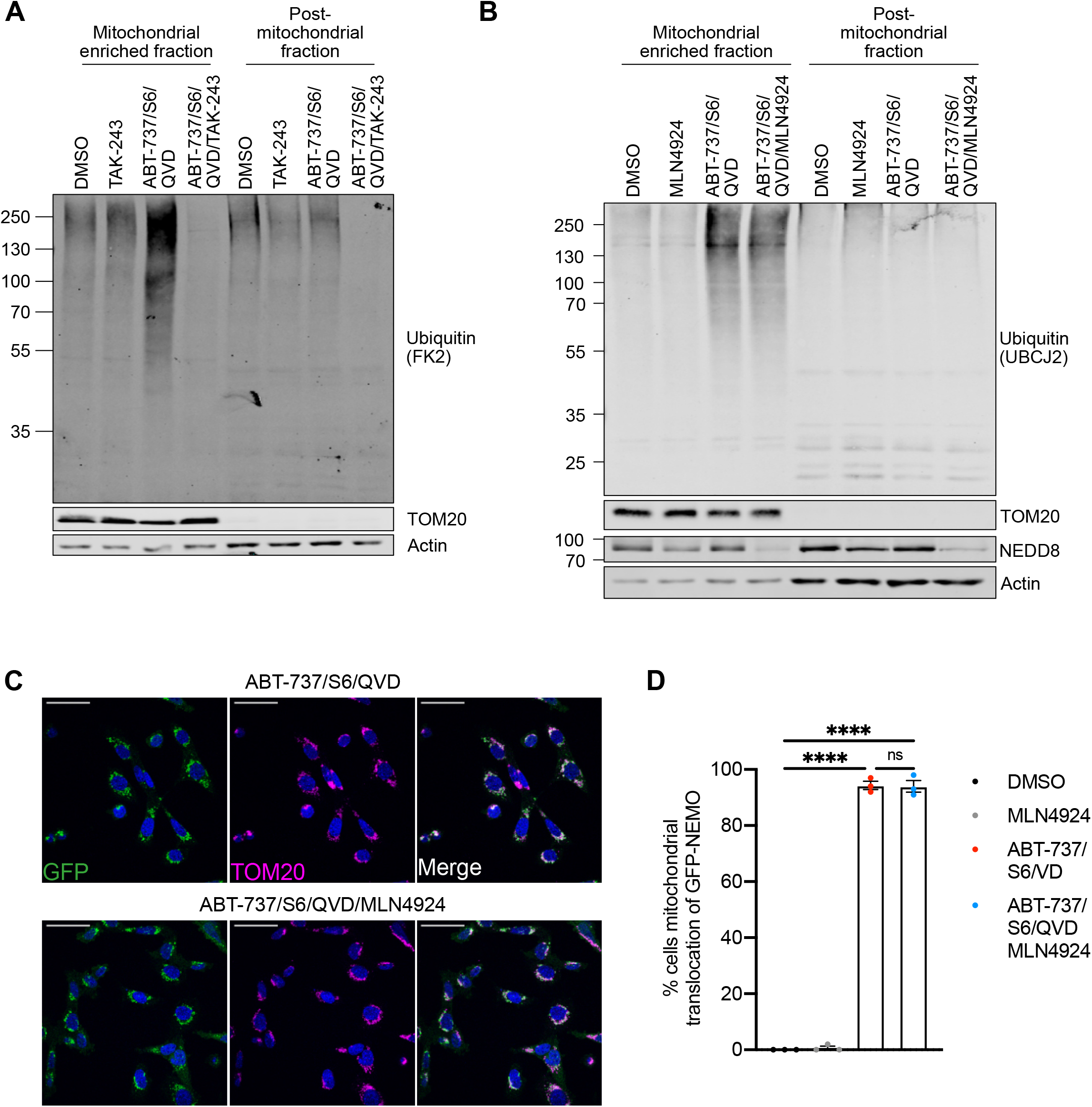
Mitochondrial ubiquitylation and GFP-NEMO translocation can be blocked by E1 inhibition and is independent of neddylation. A) SVEC4-10 cells pre-treated with 2 μΜ TAK-243 for 1 hour followed by additional 1 hour treatment with 10 μΜ ABT-737, 10 μΜ S63845 and 30 μΜ Q-VD-OPh with or without the additional of 2 μΜ TAK-243. Blots are representative for 4 independent experiments. B) SVEC4-10 cells pre-treated with 1 μΜ MLN4924 (NAE inhibitor) for 1 hour followed by additional 1 hour treatment with 10 μΜ ABT-737, 10 μΜ S63845 and 30 μΜ Q-VD-OPh with or without 1 μΜ MLN4924. Blots are representative for 2 independent experiments. C) SVEC4-10 cells expressing GFP-NEMO pre-treated with 1 μΜ MLN4924 for 1 hour followed by additional 1 hour treatment with 10 μΜ ABT-737, 10 μΜ S63845 and 30 μΜ Q-VD-OPh with or without 1 μΜ MLN4924. Cells were immunostained for TOM20 and DAPI. Images are representative for 3 independent experiments and are shown with a 50 μm scale bar. D) Quantification of C showing the percentage of cells with mitochondrial translocation of GFP-NEMO. Statistics were performed using two-way ANOVA with Tukey correction. **** p < 0.0001.

**Supplemental Figure 3.**
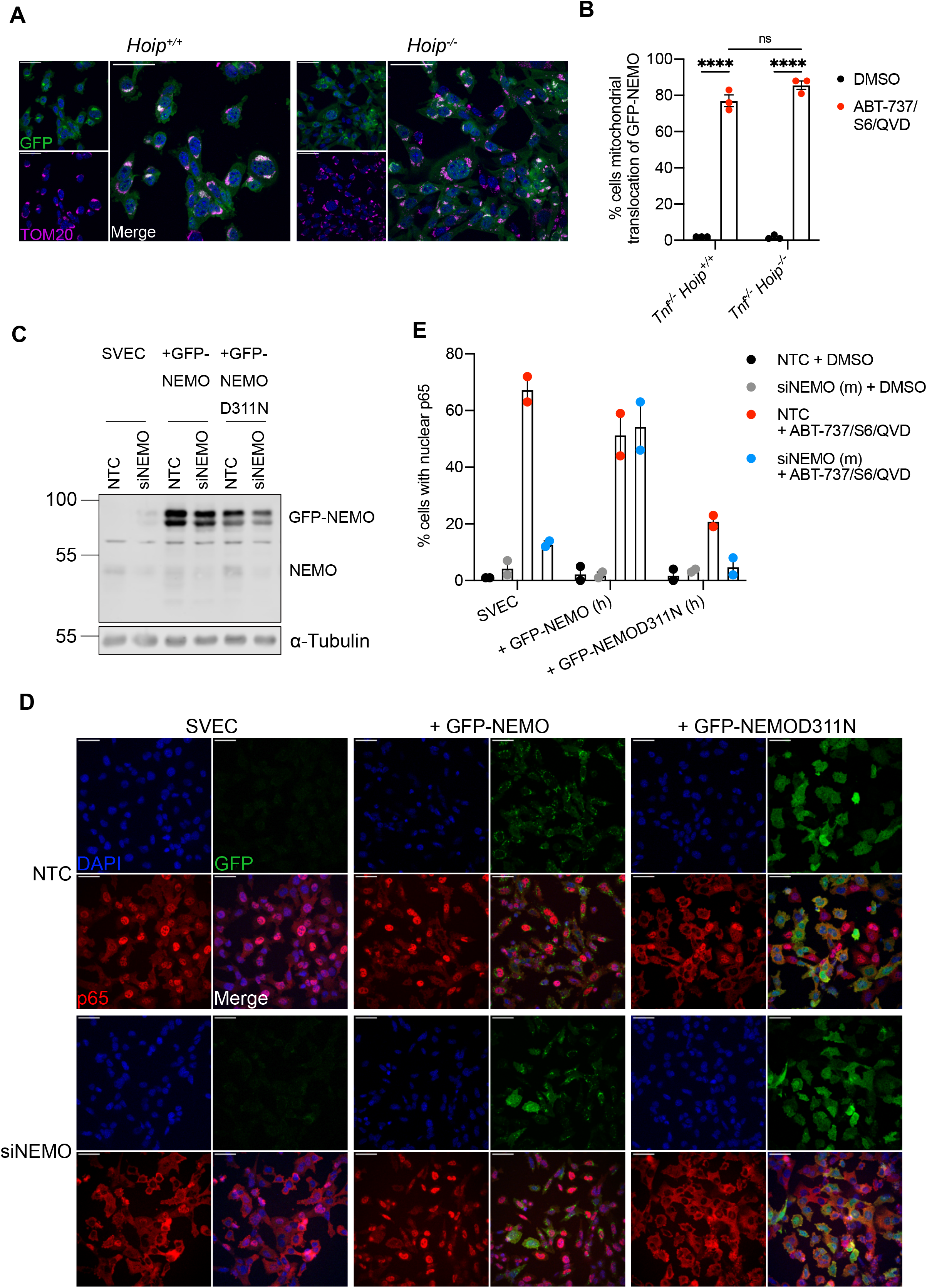
Loss of NEMO cannot be rescued during CICD by expressing non-ubiquitin binding mutants of NEMO. A) MEF *Tnf^-/-^ Hoip^+/+^* and *Tnf^-/-^ Hoip^+/+^* expressing GFP-NEMO were treated for 3 hours with 10 μΜ ABT-737, 5 μΜ S63845 and 30 μΜ Q-VD-OPh. Cells were immunostained for TOM20 and DAPI. Images are representative of 3 independent experiments. B) Quantification of A showing the percentage of cells with mitochondrial translocation of GFP-NEMO. Statistics were performed using two-way ANOVA with Tukey correction. C) Validation of SVEC4-10, SVEC4-10 GFP-NEMO and SVEC4-10 GFP-D311N cells transfected with NTC or siNEMO. Lysates were blotted for NEMO and α-tubulin. D) SVEC4-10, SVEC4-10 GFP-NEMO and SVEC4-10 GFP-D311N cells transfected with NTC or siNEMO were treated for 1 hour with 10 μΜ ABT-737, 10 μΜ S63845 and 30 μΜ Q-VD-OPh. Cells were immunostained for p65 and DAPI. Images are representative of 2 independent experiments. E) Quantification of D showing the percentage of cells with nuclear translocation of p65. **** p < 0.0001.

**Supplemental Figure 4.**
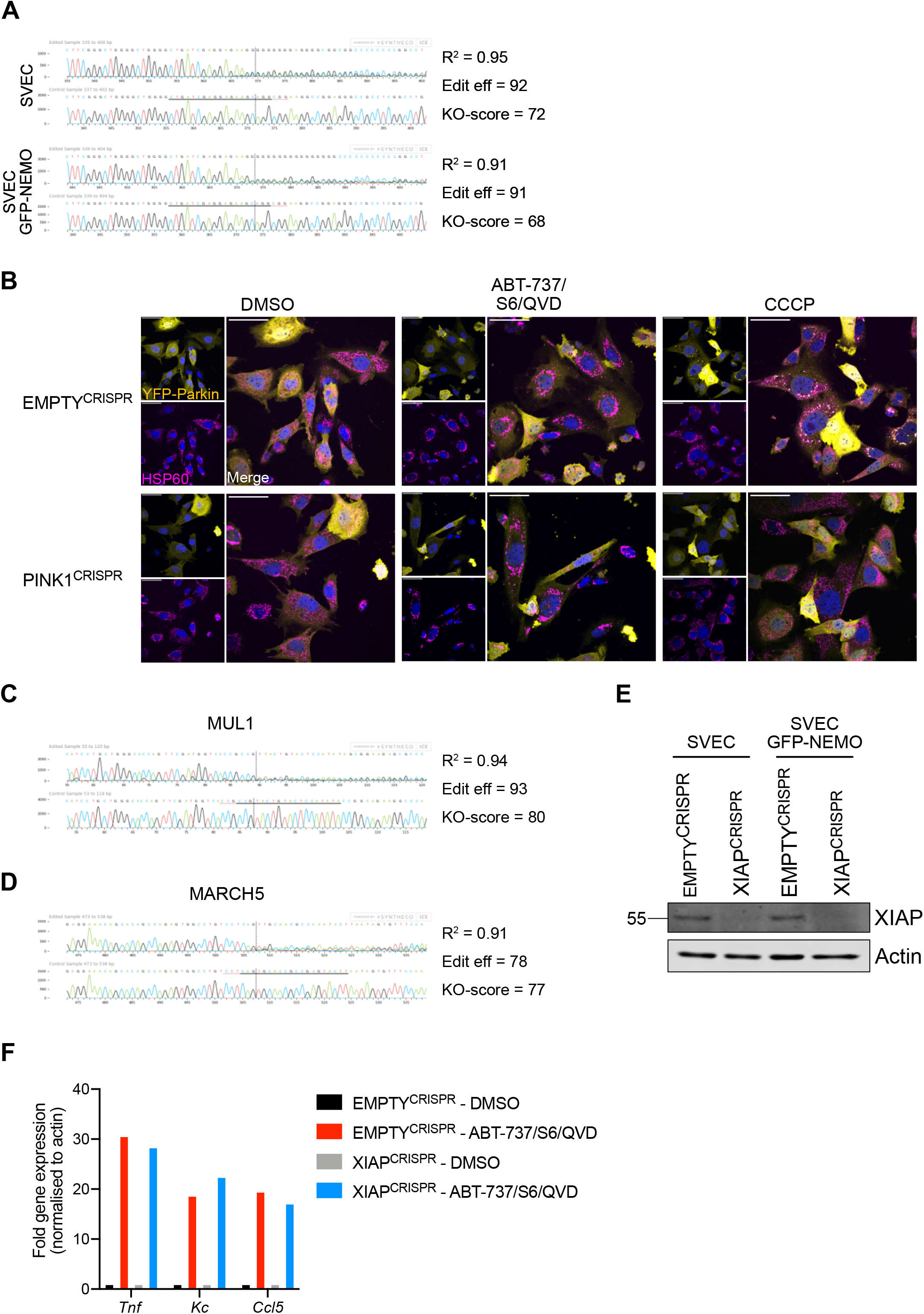
Validation of PINK1^CRISPR^, NIK^CRISPR^, MUL1MARCH5^CRISPR^ and XIAP^CRISPR^ knock-out cell lines. A) Validation of PINK1 knock-out in SVEC4-10 cells with or without GFP-NEMO expression using genomic PCR and ICE (interference of CRISPR edits) analysis. B) SVEC4-10 EMPTY^CRISPR^ and PINK1^CRISPR^ cells expressing YFP-Parkin were treated for 1 hour with 10 μΜ ABT-737, 10 μΜ S63845 and 30 μΜ QVD for 3 hours with 10 μΜ CCCP. Mitochondria were immunostained with HSP60 and DAPI. Images are representative of 2 independent experiments and displayed with 50 μm scale bar. C) Validation of MUL1 knock-out in SVEC4-10 MUL1MARCH5^CRISPR^ cells using genomic PCR and ICE analysis. D) Validation of MARCH5 knock-out in SVEC4-10 MUL1MARCH5^CRISPR^ cells using genomic PCR and ICE analysis. E) Validation of SVEC4-10 XIAP^CRISPR^ cells with and without GFP-NEMO expression using western blot. Lysates were blotted for XIAP and actin. F) *Tnf*, *Kc,* and *Ccl5* expression of SVEC4-10 EMPTY^CRISPR^ and XIAP^CRISPR^ cells treated with 10 μΜ ABT-737, 10 μΜ S63845 and 30 μΜ QVD for 3 hours. Graph is representative for 3 independent experiments.

**Supplemental Figure 5.**
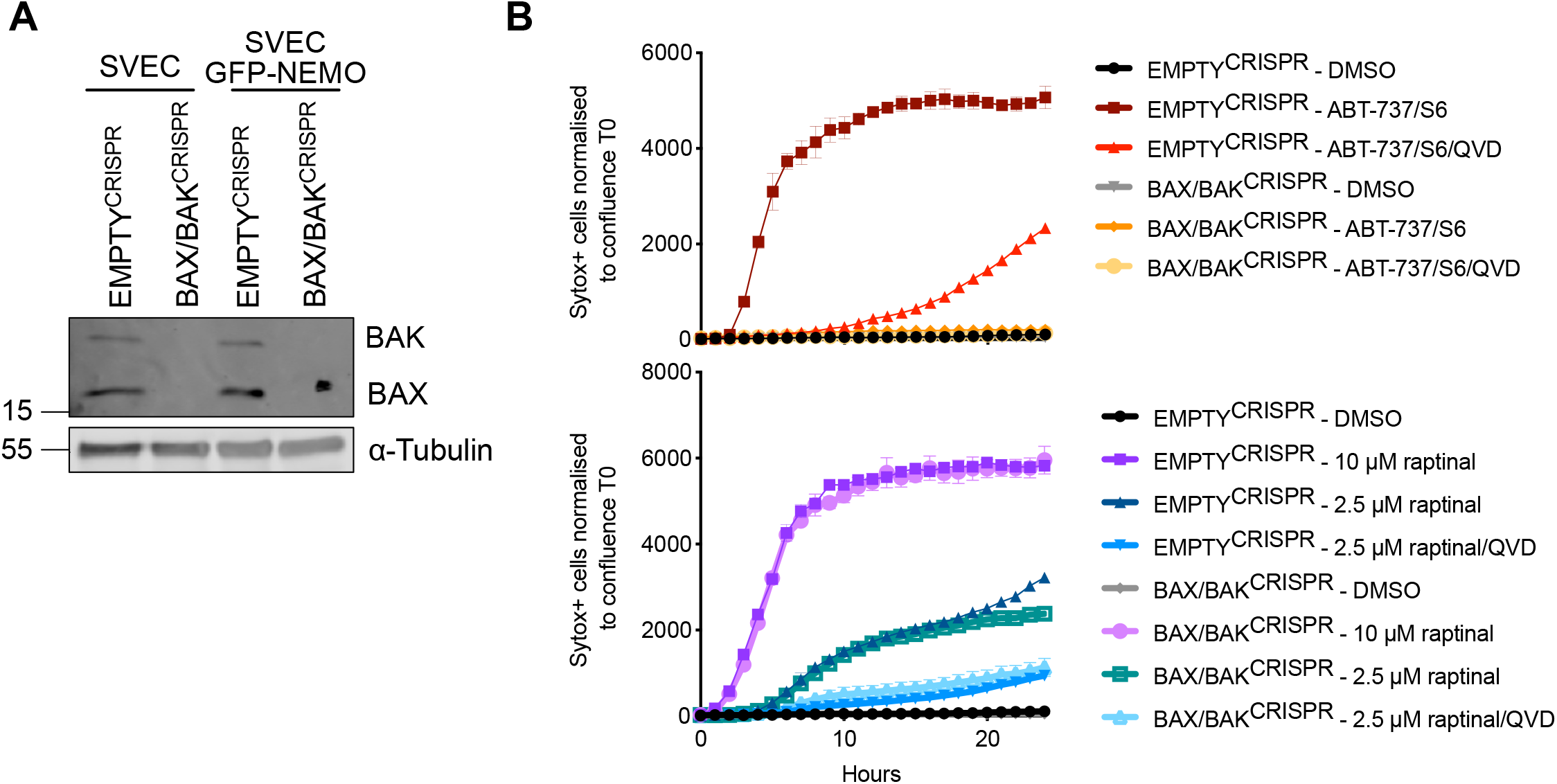
Raptinal induces cell death independent of mitochondrial permeabilization by BAX and BAK. A) EMPTY^CRISPR^ and BAX/BAK^CRISPR^ validation of SVEC4-10 cells and SVEC4-10 cells expressing GFP-NEMO. Lysates for blotted for BAX, BAK and α-tubulin. B) SVEC4-10 EMPTY^CRISPR^ and SVEC4-10 BAX/BAK^CRISPR^ cells treated with 10 μΜ ABT-737 and 10 μΜ S63845 or treated with 2.5 or 10 μΜ raptinal. Caspase-dependency of death was assessed using 30 μΜ Q-VD-OPh. Cell viability was measured using Sytox Green exclusion. Graphs are representative of 2 independent experiments and display the mean and SEM of 2 replicates. Statistics performed using two-way ANOVA with Tukey correction. *** p < 0.001, **** p < 0.0001. **Supplemental Table 1** Intramitochondrial localisation of proteins found to be ubiquitylated upon MOMP

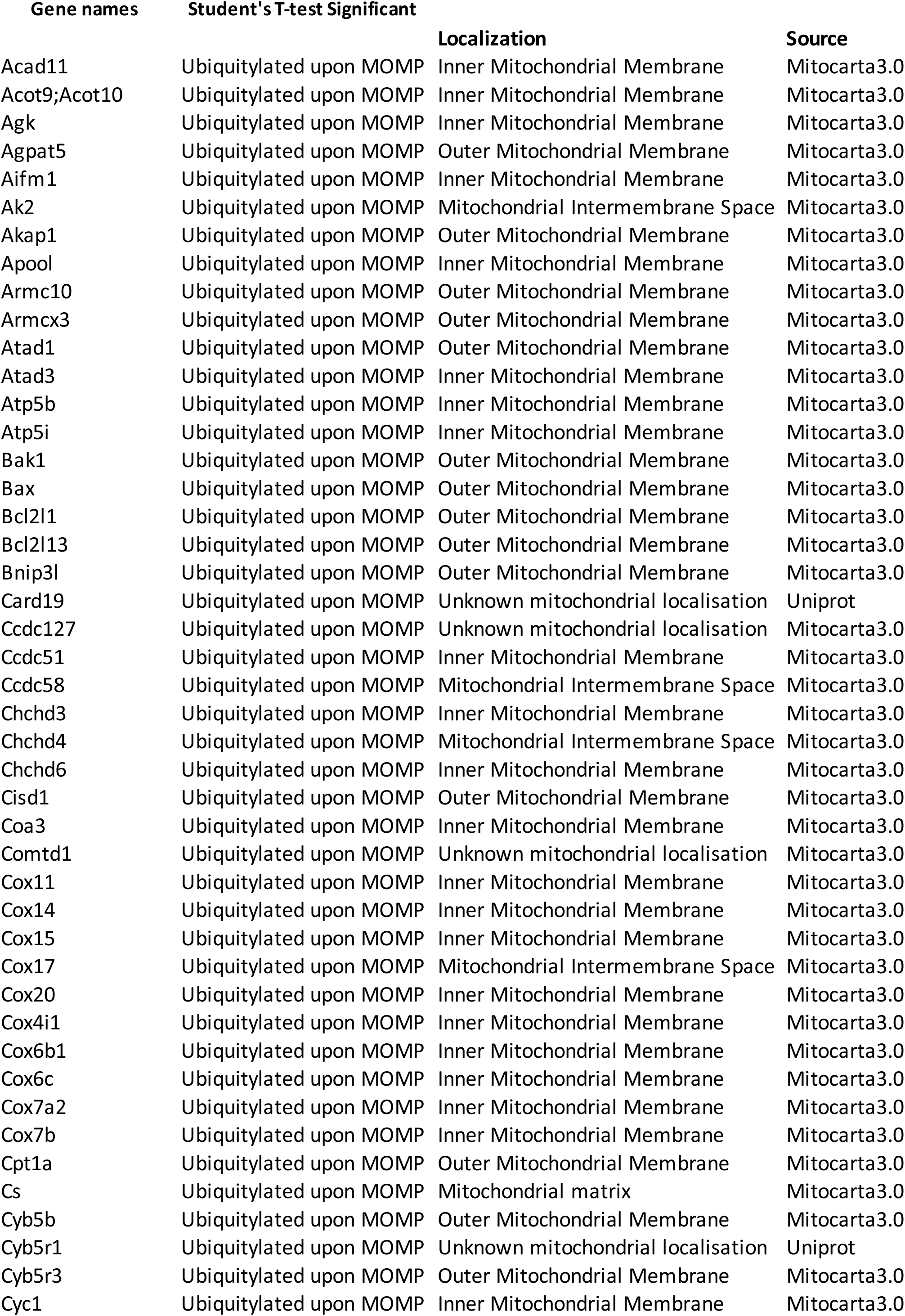
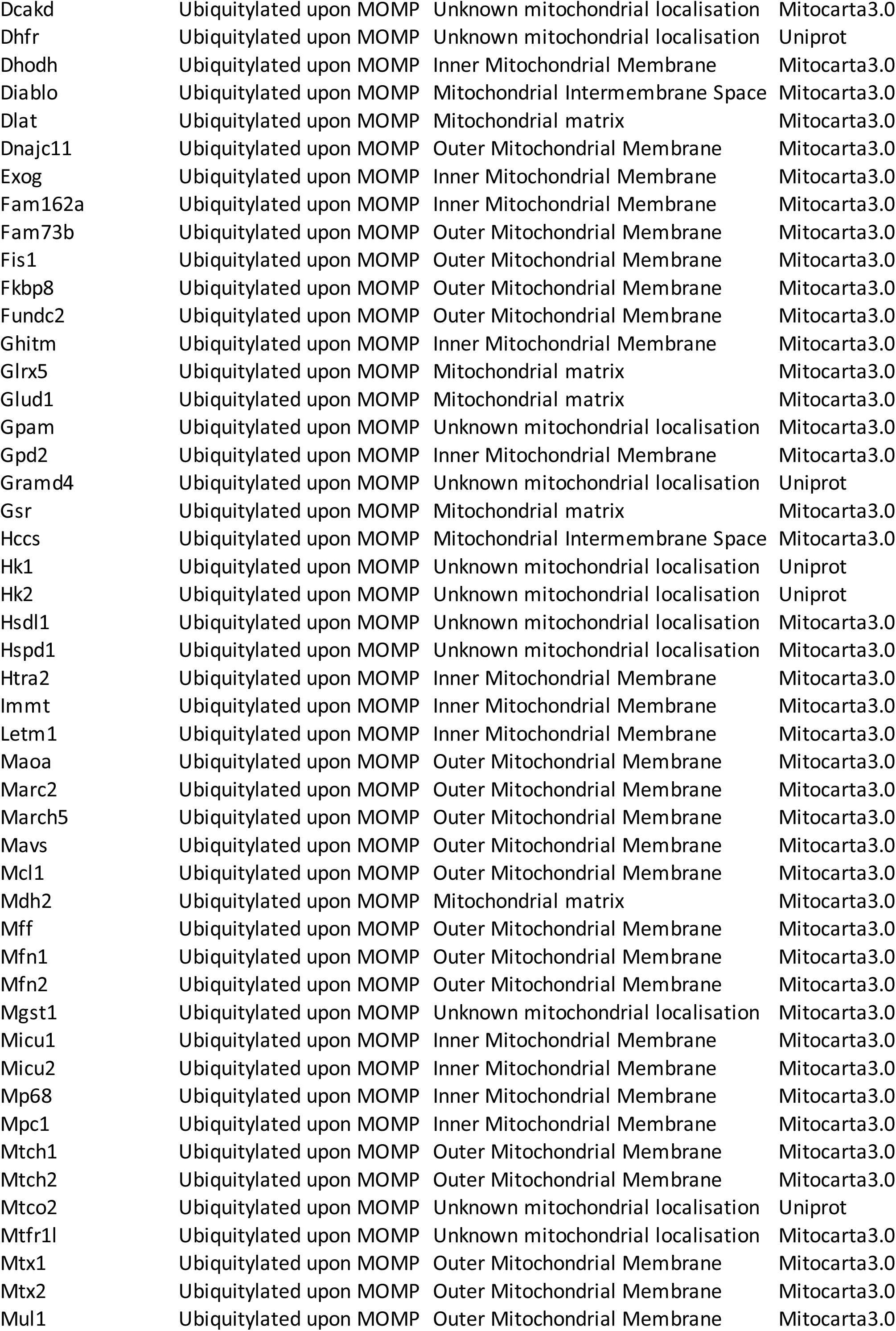
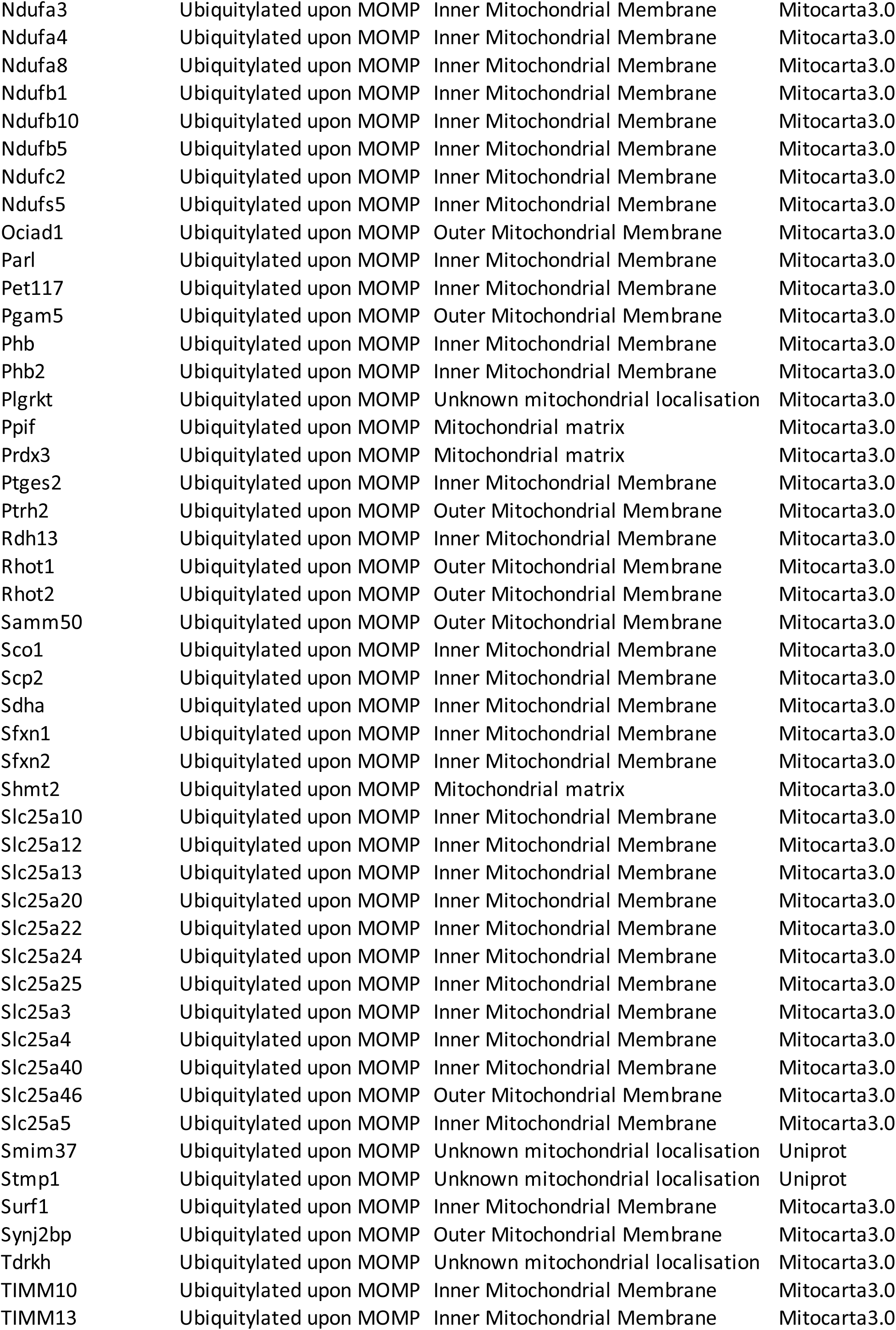
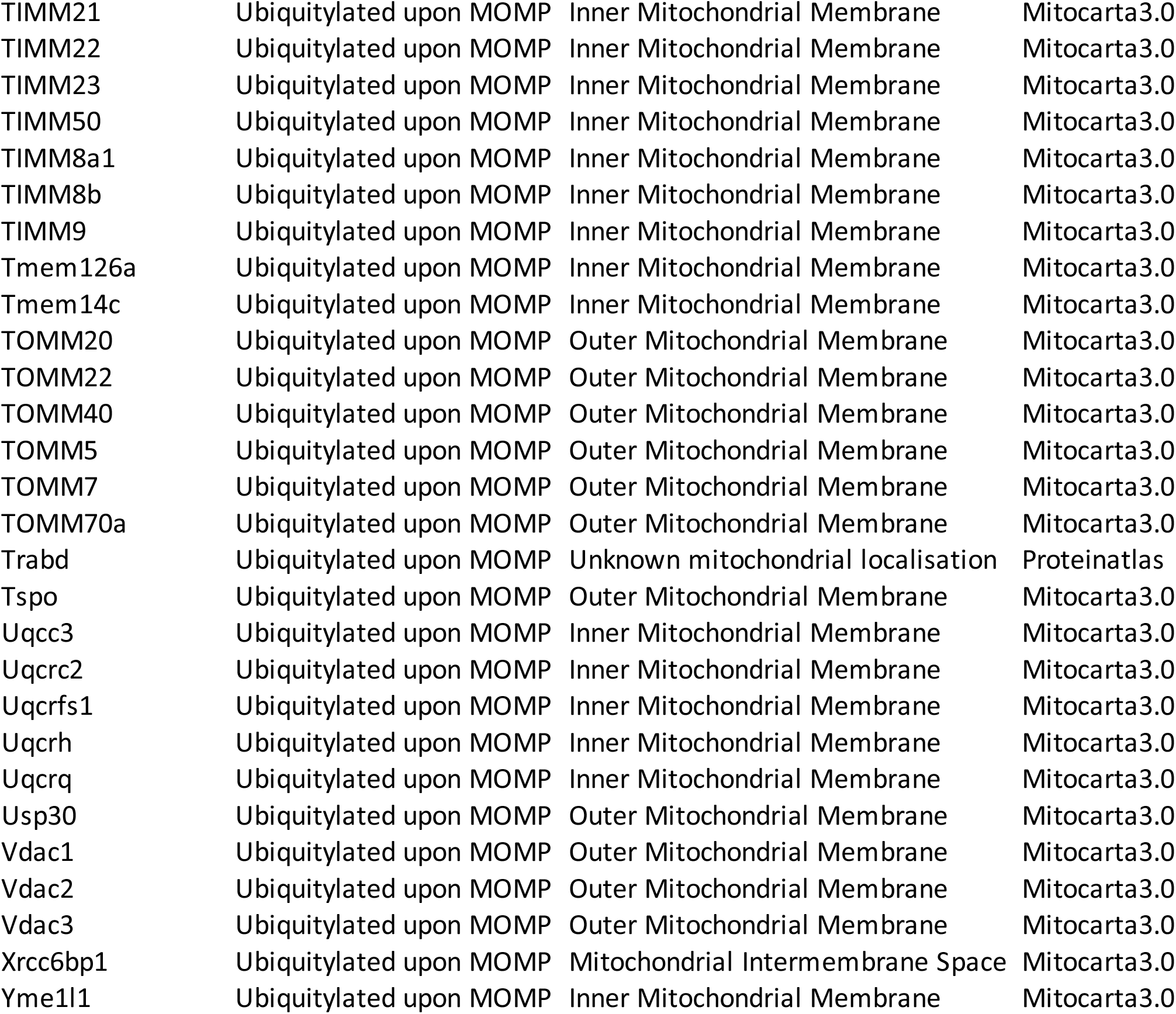

